# Deciphering *TP53* mutant Cancer Evolution with Single-Cell Multi-Omics

**DOI:** 10.1101/2022.03.28.485984

**Authors:** Alba Rodriguez-Meira, Ruggiero Norfo, Wei Xiong Wen, Agathe L. Chédeville, Haseeb Rahman, Jennifer O’Sullivan, Guanlin Wang, Eleni Louka, Warren W. Kretzschmar, Aimee Paterson, Charlotte Brierley, Jean-Edouard Martin, Caroline Demeule, Matthew Bashton, Nikolaos Sousos, Angela Hamblin, Helene Guermouche, Florence Pasquier, Christophe Marzac, François Girodon, Mark Drummond, Claire Harrison, Isabelle Plo, Sten Eirik W. Jacobsen, Bethan Psaila, Supat Thongjuea, Iléana Antony-Debré, Adam J Mead

**Affiliations:** Haematopoietic Stem Cell Biology Laboratory, Medical Research Council Molecular Haematology Unit, Medical Research Council Weatherall Institute of Molecular Medicine, University of Oxford, Oxford OX3 9DS, UK; NIHR Biomedical Research Centre, University of Oxford, Oxford, OX3 9DS, United Kingdom; Medical Research Council Centre for Computational Biology, Weatherall Institute of Molecular Medicine, University of Oxford, Oxford OX3 9DS, UK; INSERM, UMR 1287, Villejuif, France; Gustave Roussy, Villejuif, France; Université Paris Saclay, Gif-sur-Yvette, France; Université de Paris, Paris, France; Department of Cell and Molecular Biology, Karolinska Institutet, SE-171 77 Stockholm, Sweden; Karolinska University Hospital, Stockholm, Sweden; Center for Hematology and Regenerative Medicine, Department of Medicine Huddinge, Karolinska Institutet, Karolinska University Hospital, SE-141 86 Stockholm, Sweden; Center for Hematological Malignancies, Memorial Sloan Kettering Cancer Center, New York, NY, USA; Laboratoire d’Hématologie, CHU Dijon, Dijon, France; The Hub for Biotechnology in the Built Environment, Faculty of Health and Life Sciences, Northumbria University, Newcastle upon Tyne, NE1 8S, UK; National Institute for Health Research Biomedical Research Centre, University of Oxford, Oxford, UK; Sorbonne Université, INSERM, Centre de Recherche Saint-Antoine, AP-HP, Hôpital Saint-Antoine, Service d’hématologie biologique, F-75012, Paris; Département d’Hématologie, Gustave Roussy, Villejuif, France; Laboratoire d’Immuno-Hématologie, Gustave Roussy, Villejuif, France; INSERM, UMR 866, Centre de Recherche, Dijon, France; Beatson Cancer Centre, Glasgow, UK; Guy’s and St Thomas’ NHS Foundation Trust, Department of Haematology, London, UK

**Author notes:** These authors contributed equally (Ruggiero Norfo, Wei Xiong Wen, Agathe L. Chédeville, Haseeb Rahman). These authors contributed equally (Iléana Antony-Debré, Adam J Mead).

## Abstract

*TP53* is the most commonly mutated gene in human cancer, typically occurring in association with complex cytogenetics and dismal outcomes. Understanding the genetic and non-genetic determinants of *TP53-*mutation driven clonal evolution and subsequent transformation is a crucial step towards the design of rational therapeutic strategies. Here, we carry out allelic resolution single-cell multi-omic analysis of haematopoietic stem/progenitor cells (HSPC) from patients with a myeloproliferative neoplasm who transform to *TP53-*mutant secondary acute myeloid leukaemia (AML), a tractable model of *TP53*-mutant cancer evolution. All patients showed dominant *TP53 ‘*multi-hit’ HSPC clones at transformation, with a leukaemia stem cell transcriptional signature strongly predictive of adverse outcome in independent cohorts, across both *TP53-*mutant and wild-type AML. Through analysis of serial samples and antecedent *TP53*-heterozygous clones, we demonstrate a hitherto unrecognised effect of chronic inflammation, which supressed *TP53* wild-type HSPC whilst enhancing the fitness advantage of *TP53* mutant cells. Our findings will facilitate the development of risk-stratification, early detection and treatment strategies for *TP53*-mutant leukaemia, and are of broader relevance to other cancer types.

## Main Text

Tumour protein 53 (*TP53*) is the most frequently mutated gene in human cancer, typically occurring as a multi-hit process with point mutation of one allele and loss of the other wild-type allele^1, 2^. *TP53* mutations are also strongly associated with copy number alterations (CNA) and structural variants, reflecting the role of p53 in the maintenance of genomic integrity^2, 3^. In myeloid malignancies, presence of a *TP53* mutation defines a distinct clinical entity^1^, associated with complex CNA, lack of response to conventional therapy and dismal outcomes^2, 4, 5^. Understanding the mechanisms by which TP53 mutations drive clonal evolution and disease progression is a crucial step towards the development of rational strategies to diagnose, stratify, treat and potentially prevent this condition.

Myeloproliferative neoplasms (MPN) arise in haematopoietic stem cells (HSC) through the acquisition of mutations in JAK/STAT signalling pathway genes (*JAK2*, *CALR* or *MPL),* leading to aberrant proliferation of myeloid lineages^6^. Progression to secondary acute myeloid leukaemia (sAML) occurs in 10-20% of MPN and is characterized by cytopenias, increased myeloid blasts, acquisition of aberrant leukaemia stem cell (LSC) properties by haematopoietic stem/progenitor cells (HSPC) and median survival of less than one year^7, 8^. *TP53* mutations are detected in approximately 20-35% of post-MPN sAML^9–11^ (collectively termed *TP53*-sAML), often in association with loss of the remaining wild-type allele^12^ and multiple CNAs^13^. Furthermore, deletion of *Trp53* combined with *JAK2V617F* mutation leads to a highly penetrant myeloid leukaemia in mice^11, 14^.

Notwithstanding the established role of *TP53* mutation in MPN transformation, *TP53*-mutant subclones are also present in 16% of chronic phase MPN (CP-MPN) and in most cases this does not herald the development of *TP53*-sAML^15^. However, little is known about the additional genetic and non-genetic determinants of clonal evolution following the acquisition of a *TP53* mutation. Resolving this question requires multiple layers of intratumoural heterogeneity to be unravelled, including reliable identification of the *TP53* mutation, loss of the wild-type allele and presence of CNA. Integrating this mutational landscape with cellular phenotype and transcriptional signatures will resolve aberrant haematopoietic differentiation and molecular properties of LSC in *TP53*-sAML. This collectively requires single-cell approaches which combine molecular and phenotypic analysis of HSPCs with allelic-resolution mutation detection, an approach recently enabled by the TARGET-seq technology^16^.

### Convergent clonal evolution during *TP53-*driven leukaemic transformation

To characterize the genetic landscape of *TP53*-sAML, we analysed 33 *TP53*-sAML patients (TableS1) through bulk-level targeted next generation sequencing and SNP array (Extended Data Fig.1). We detected MPN-driver mutations (*JAK2*, *CALR*) in 28 patients (85%), and co-occurring myeloid driver mutations in 24 patients (73%). Multiple *TP53* mutations were present in one third (n=11) of patients, including 2 patients with 3 *TP53* mutations. 82% (18/22) of patients with a single *TP53* mutation showed a high variant allelic frequency (VAF) of >50%. CNAs were present in all patients analysed, and 87% (20/23) had a complex karyotype (≥ 3 CNA; Extended Data Fig.1a-g). Deletion or copy neutral loss of heterozygosity affecting the *TP53* locus (chr17p13.1) was detectable at the bulk level in 43% of patients (10/23) (Extended Data Fig.1b-d). Taken together, these findings support that *TP53-*sAML is associated with complex genetic intratumoural heterogeneity.

To characterize tumour phylogenies and subclonal structures, we performed TARGET-seq analysis^16^ on 17608 Lin^-^CD34^+^ HSPCs from 14 *TP53*-sAML patients (Extended Data Fig.1a), 9 age-matched healthy donors (HD) and 8 previously published myelofibrosis (MF) patients (Fig.1a, gating strategy shown in Extended Data Fig.2a). HSPCs wild-type for all mutations were present in 10 of 14 patients (Extended Data Fig.2b-o), providing a valuable population of cells for intra-patient comparison with mutation-positive cells^17^. In all cases, the dominant clone showed loss of wild-type *TP53* through 4 patterns of clonal evolution: (1) biallelic *TP53* mutations by acquisition of a second mutation on the other *TP53* allele, (2) hemizygous *TP53* mutations (deleted *TP53* wild-type allele), (3) parallel evolution with 2 clones harbouring different *TP53* alterations, (4) a *JAK2* negative dominant clone with biallelic *TP53* mutations in patients with previous *JAK2*-mutant MPN^18^ (Fig.1b-e, Extended Data Fig.2b-o). Biallelic mutations were confirmed by single molecule cloning and computational analysis (Extended Data Fig.1h-j). Integration of index-sorting data revealed that dominant *TP53* multi-hit clones were enriched in progenitor populations as previously described in *de novo* AML^19^, whereas *TP53*-mutant cells were rare in the HSC (Lin^-^CD34^+^CD38^-^CD45RA^-^CD90^+^) compartment (Extended Data Fig.3a). CNA analysis using single-cell transcriptomes showed that all *TP53* multi-hit clones harboured at least one highly clonally-dominant CNA, with very few *TP53-*mutant cells without evidence of a CNA (3.4±1.2%) and an additional 5/14 (36%) patients also showing cytogenetically-distinct subclones (Fig.1f,g, Extended Data Fig.2p,q).

**Fig. 1.**
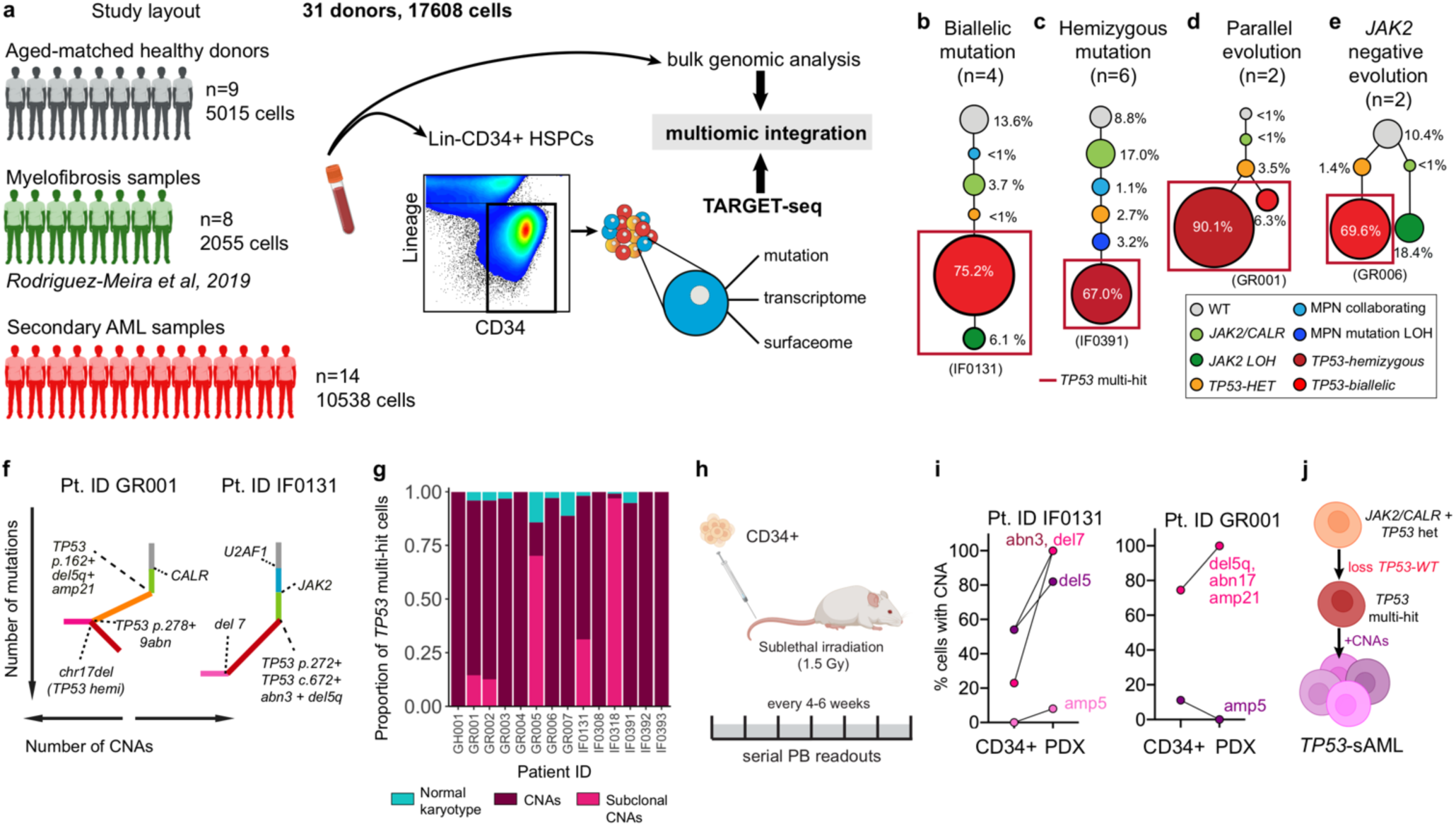
Clonal evolution of *TP53*-sAML. **a,** Schematic study layout for TARGET-seq profiling of 17608 Lin^-^ CD34^+^ HSPCs from 31 donors. **b-e,** Representative examples of the four major patterns of clonal evolution in *TP53*-sAML patients: bi-allelic mutations **(b),** hemizygous mutations **(c)**, parallel evolution **(d)** and *JAK2* negative biallelic evolution **(e)**. The numbers in parenthesis indicate the number of patients in each category. The size of the circles is proportional to each clone’s size, indicated as a percentage of total Lin^-^CD34^+^ cells for one representative patient in each group; each clone is coloured according to its genotype (Related to Extended Data Fig.2b-o) and red boxes indicate *TP53* multi-hit clones. **f,** Representative examples from integrated mutation and CNA-based clonal hierarchies. Solid lines indicate acquisition of a genetic hit (i.e. point mutation or CNA) whereas dotted lines indicate the specific genetic hit acquired in each step of the hierarchy (Related to Extended Data Fig.2p,q). **g,** Proportion of *TP53* multi-hit cells classified as carrying clonal or subclonal CNAs in each patient, using a transcriptomic-based CNA clustering approach (inferCNV). **h,** Experimental strategy for xenotransplantation of CD34^+^ cells from *TP53*-sAML patients in immunodeficient mice. **i,** Percentage of cells carrying CNAs found in each PDX and corresponding Lin^-^ CD34^+^ cells from the primary *TP53*-sAML sample transplanted (Related to Supplementary Fig.3). **j,** Model of *TP53*-sAML genetic evolution.

To confirm that dominant HSPC clones were functional LSCs, we established patient-derived xenografts (PDX) for 2 *TP53*-sAML patients (Fig.1h). Mice developed leukaemia in 27-31 weeks with high numbers of human CD34^+^ myeloid blast cells in the bone marrow (BM) (Extended Data Fig.3b-d), with a progenitor phenotype, *TP53* mutations and CNAs similarly to the dominant clone from patients’ primary cells (Fig. 1i, Extended Data Fig.3e-l). In Patient IF0131, a monosomy 7 subclone (Fig.1f) preferentially expanded in PDX models (Fig.1i). Monosomy 7 cells showed a distinct transcriptional profile with increased WNT, RAS, MAPK signalling and cell cycle associated transcription (Extended Data Fig.3m,n). Together, these data are compatible with a fitness advantage of monosomy 7 cells, a recurrent event in *TP53-*sAML (Extended Data Fig.1b,c), driven by activation of signalling pathways which may relate to deletion of chromosome 7 genes such as *EZH2*^20^. In summary, the dominant leukaemic clones in *TP53-*sAML were invariably characterized by multiple hits affecting *TP53* (“multi-hit” state), indicating strong selective pressure for complete loss of wild-type *TP53*, together with gain of CNAs and complex cytogenetic evolution, with very few *TP53* multi-hit cells with a normal karyotype (Fig.1j).

### Molecular signatures of *TP53*-mutant mediated transformation

To understand the cellular and molecular framework through which *TP53* mutation drives clonal evolution, we next analysed single-cell RNA-seq data from 10538 *TP53-*sAML HSPCs alongside 2055 MF and 5015 HD HSPCs passing quality control. Force-directed graph analysis revealed separate clustering of *TP53*-mutant HSPC in comparison with HD and MF cells, with a high degree of inter-patient heterogeneity (Extended Data Fig.4a) as observed in other haematopoietic malignancies^21^. This could potentially be explained by patient-specific cooperating mutations and cytogenetic alterations (Extended Data Fig.1). TARGET-seq analysis uniquely enabled comparison of *TP53* multi-hit HSPC to *TP53* wild-type preleukaemic stem cells (“preLSCs”) from the same *TP53*-sAML patients as well as HD and MF, to derive a specific *TP53* multi-hit signature including known p53-pathway genes (Extended Data Fig.4b, Table S2).

Integration of single cell transcriptomes and diffusion map analysis of HSPC from *TP53-*sAML patients showed that *TP53* multi-hit HSPC clustered separately from *TP53* wild-type preLSC in two distinct populations with enrichment of LSC and erythroid-associated transcription respectively (Fig.2a, Table S3), and a differentiation trajectory towards the erythroid-biased population (Fig.2b), an unexpected finding given that erythroleukaemia is uncommon in *TP53-*sAML^22, 23^. *TP53* multi-hit LSCs showed enrichment of cell cycle, inflammatory, signalling pathways and LSC associated transcription, whereas *TP53* multi-hit erythroid cells were depleted of the latter (Extended Data Fig.4c).

**Fig. 2.**
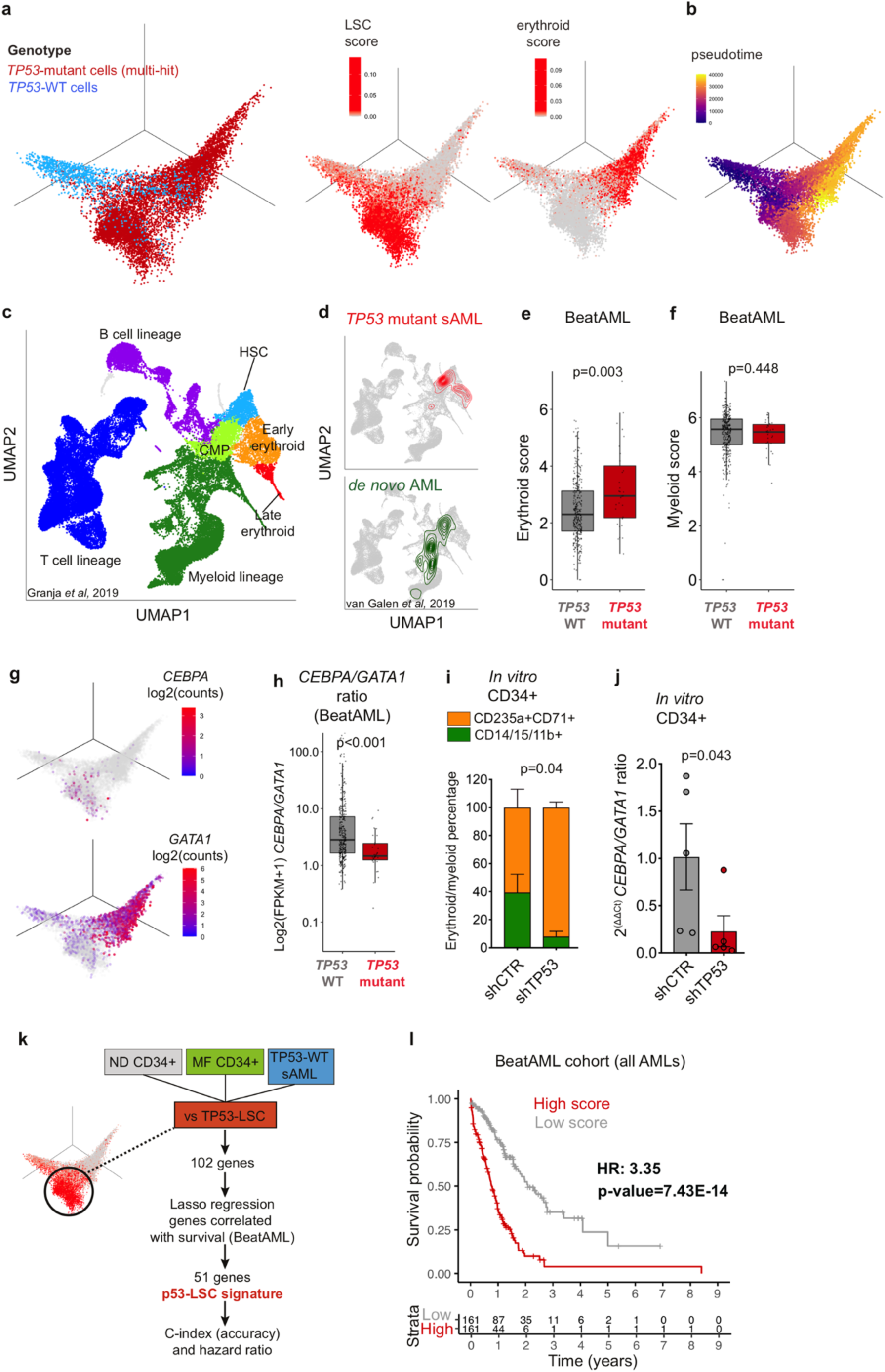
Distinct differentiation trajectories and molecular features of *TP53*-sAML. **a,** Three dimensional diffusion map of 10539 Lin^-^CD34^+^ cells from 14 sAML samples coloured by *TP53* genotype (left), leukemic stem cell score (middle) and erythroid transcription score (right). **b,** Monocle3 pseudotime ordering of the same single cells as in **(a)**. **c-d,** UMAP representation of a healthy donor hematopoietic hierarchy (**c**; Granja *et al*, 2019) and latent semantic index projection of *TP53* multi-hit cells from 14 sAML patients **(d, top)** and cells from *de novo* AML patients **(d, bottom;** van Galen *et al*, 2019) onto the healthy donor hematopoietic hierarchy atlas **(c)**. **e-f,** Expression of an erythroid **(e)** and myeloid **(f)** gene score in AML patients from the BeatAML dataset stratified by *TP53* mutational status (n=329 *TP53-*WT; n=31 *TP53-*mutant). Boxplots indicate median and quartiles; “p” indicates Wilcoxon rank sum test p-value. **g,** *CEBPA* (top) and *GATA1* (bottom) expression in the same cells as in (a-b). **h,** *CEBPA* and *GATA1* expression ratio in the same patient cohort as in (e,f). **i-j,** Proportion of immature erythroid (CD235a^+^CD71^+^) and myeloid (CD14^+^, CD15^+^ or CD11b^+^) cells (**i**) and ratio of *CEBPA* to *GATA1* expression in total cells **(j)** after 12 days of differentiation of peripheral blood CD34^+^ cells from patients with MPN transduced with shRNA targeting *TP53* or a scramble control (shCTR). n=5 patients, 3 independent experiments. Barplot indicates mean ± s.e.m. and “p”, two-tailed paired t-test p-value. (Related to Extended Data Fig.5i). **k,** Schematic representation of the key analytical steps to derive a 51-gene *TP53*-LSC sAML signature**. l,** Kaplan-Meier analysis of AML patients (n=322) from the BeatAML cohort stratified by p53-LSC signature score (high: above median; low, below median) derived in **(k)** (Related to Supplementary Fig.6). HR: hazard ratio. “p” indicates log-rank test p-value.

To further explore this erythroid-biased population, we projected *TP53* multi-hit cells onto a previously published healthy donor haematopoietic hierarchy^24^. *TP53*-sAML differed from *de novo* AML with an enrichment into HSC and early erythroid populations, whereas *de novo* AML were enriched in myeloid progenitors (Fig.2c,d)^25^. A similar enrichment was observed for *TP53* multi-hit cells when mapped on a Lin^-^CD34^+^ MF cellular hierarchy (Extended Data Fig.5a,b), supporting an aberrant erythroid-biased differentiation trajectory in *TP53*-sAML.

To determine whether upregulation of erythroid-associated transcription was a more widespread phenomenon in *TP53-*mutant AML, we investigated erythroid-myeloid associated transcription in the BeatAML and TCGA cohorts^26, 27^. Erythroid scores were increased in *TP53* mutant compared to *TP53* wild-type AML, whereas there was no significant difference in myeloid scores (Fig.2e-f, Extended Data Fig.5c-f, scores described in Table S3). We next investigated the expression of key transcription factors for erythroid/granulomonocytic commitment and found increased *GATA1* expression in Lin^-^CD34^+^ *TP53* multi-hit HSPC, whereas *CEBPA* was only expressed at low levels (Fig.2g). Analysis of the BeatAML cohort revealed increased *GATA1* and reduced *CEBPA* expression in association with *TP53* mutation (Extended Data Fig.5g), with consequent reduction in the *CEBPA/GATA1* expression ratio (Fig.2h). Similar findings were observed in *TP53* knock-out or mutant isogenic MOLM13 cell lines (Extended Data Fig.5h)^28^. These observations suggest that the *CEBPA/GATA1* expression ratio, an important transcription factor balance which affects erythroid versus myeloid differentiation in leukaemia^29, 30^ is disrupted by *TP53* mutation.

To determine whether p53 directly influences myeloid-erythroid differentiation, we knocked-down *TP53* in *JAK2*V617F CD34^+^ cells from MPN patients (Extended Data Fig.5i). *TP53* knock-down led to increased erythroid (CD71^+^CD235a^+^) and decreased myeloid (CD14^+^/CD15^+^/CD11b^+^) differentiation *in vitro* (Fig.2i) and consequently decreased *CEBPA*/*GATA1* expression ratio (Fig.2j), suggesting that p53 may directly contribute to the aberrant myelo-erythroid differentiation observed.

As ‘stemness scores’ have previously been applied to determine prognosis in AML^31^, we next asked whether a single-cell defined *TP53* multi-hit LSC signature might identify AML patients with adverse outcomes. Single cell multi-omics allowed us to derive a 51-gene “p53LSC-signature” (Table S4) by comparing gene expression of HD, *JAK2*-mutant MF HSPC and *TP53* wild-type preLSC to transcriptionally-defined *TP53*-mutant LSCs (Fig.2a,k). High p53LSC-signature score was strongly associated with poor survival in the independent BeatAML and TCGA cohorts, irrespective of *TP53* mutational status (Fig.2l, Extended Data Fig.6a-c). The p53LSC signature performed well as a predictor of survival, including in sAML patients, as compared to the previously published LSC17 score^31^ and p53-mutant score generated using all *TP53*-mutant HSPC (Extended Data Fig.6d,e, TableS4), providing a powerful new tool to aid risk stratification in AML.

### Preleukaemic *TP53*-wild-type cells display self-renewal and differentiation defects

TARGET-seq uniquely enabled phenotypic and molecular characterization of *TP53* wild-type preLSC obtained in sufficient numbers (>20 cells) from 9 of 14 *TP53*-sAML patients (Fig.3a). Some of these preLSC represented the antecedent CP-MPN clone with MPN-associated mutation (532/880 cells, 60.5%), whereas others were wild-type for all mutations (348/880 cells, 39.5%). PreLSC were enriched in HSC-associated genes, and superimposed on HSC clusters in HD and MF haematopoietic hierarchies (Fig.3a,b). Index sorting revealed that preLSCs were strikingly enriched in the phenotypic HSC compartment, unlike *TP53* multi-hit HSPC (Fig.3c). Pre-LSCs were rare, as reflected by a reduction in the numbers of phenotypic HSCs present within the Lin^-^CD34^+^ HSPC compartment in *TP53-*sAML compared to HD (Extended Data Fig.7a).

**Fig. 3.**
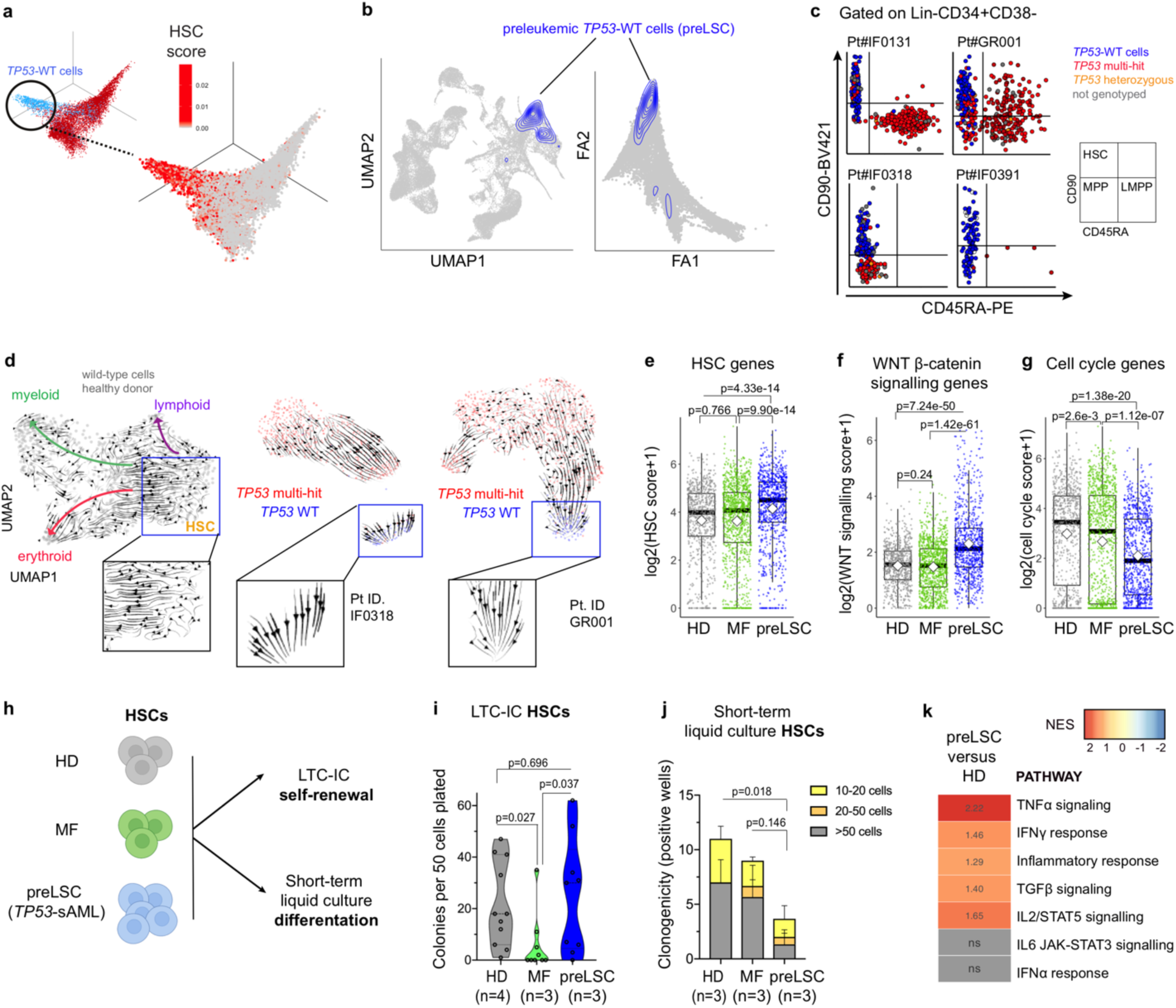
Molecular and functional analysis of preleukemic stem cells in *TP53*-sAML patients. **a,** Three dimensional diffusion map of 10539 Lin^-^CD34^+^ cells from *TP53*-sAML patients (Related to Fig.2a) coloured by expression of a HSC signature (Table S3). **b,** Projection of *TP53*-WT (n=880) preleukemic stem cells (preLSC) on a healthy donor (left) and myelofibrosis (right) hematopoietic hierarchy (related to Fig.2c, Extended Data Fig.5a). **c,** Immunophenotype of Lin^-^CD34^+^CD38^-^ cells from four representative sAML patients coloured by genotype. Lin^-^CD34^+^CD38^-^CD90^+^CD45RA^-^ cells (HSCs) were enriched using the sorting strategy outlined in Extended Data Fig.2a. **d,** scVelo analysis of differentiation trajectories of Lin^-^ CD34^+^ cells from one healthy donor (left) and two representative *TP53*-sAML patients (middle and right). Insets show HSC or preLSCs clusters. **e-g,** Scores of HSC **(e)**, WNT ß-catenin signalling **(f)** and cell-cycle **(g)** associated transcription in Lin^-^CD34^+^CD38^-^ cells from HD (n=730 cells), MF (n=1106 cells) and preLSCs from *TP53*-sAML patients (n=880 cells). Boxplot represents median and quartiles; white square indicates the mean for each group. “p” indicates Wilcoxon rank sum test p-value. **h-j,** Functional analysis of preLSC. Schematic representation of HSC *in vitro* assays (**h**), long-term culture initiating-cell (LTC-IC) colony forming unit activity **(i)** and short-term *in vitro* liquid culture clonogenicity **(j)** of HSC from HD (n=3-4), MF (n=3) and preLSCs from *TP53*-sAML patients (n=3, samples used (IF0131, IF0391, GR001) were known to have *TP53*-WT preLSC in the HSC compartment). Violin plot indicates points’ density; dashed lines, median and quartiles, 2 independent experiments **(i)**; barplot indicates mean ± s.e.m., 3 independent experiments **(j).** “p” indicates two-tailed Student t-test p-value. **k**, GSEA analysis of HALLMARK inflammatory pathways in preLSCs compared to HD; positive NES in the heatmap represents significant (FDR q-value<0.25) enrichment in preLSCs, values indicate NES for each pathway. NES: Normalized Enrichment Score. FDR: False Discovery Rate.

We reasoned that the HSC phenotype of preLSCs, with reduced frequency in progenitor compartments, might reflect impaired differentiation. To explore this hypothesis, we carried out scVelo analysis, which showed absence of a transcriptional differentiation trajectory in preLSCs, unlike HD HSCs (Fig.3d). PreLSC showed increased expression of haematopoietic stem cell and Wnt β-catenin genes and decreased cell cycle genes as compared to HD and MF cells (Fig.3e-g, TableS3). To functionally confirm these findings, we sorted phenotypic HSC (to purify preLSC), as well as other progenitor cells, from HD, MF and *TP53-*sAML patients for long term culture initiating cell (LTC-IC) and short-term cultures (Fig.3h; Extended Data Fig.7b). PreLSC LTC-IC activity was similar to HD and increased compared to MF, with preserved terminal differentiation capacity and confirmed *TP53* wild-type genotype (Fig.3i, Extended Data Fig.7c-f). In short-term liquid culture, preLSCs showed reduced clonogenicity, with retained CD34 expression and decreased proliferation (Fig.3j, Extended Data Fig.7g-h). In summary, we identified rare and phenotypically distinct preLSCs from *TP53*-sAML samples which were characterized by differentiation defects and distinct stemness, self-renewal and quiescence signatures. As these cells were *TP53*-wild-type, and showed normal differentiation after prolonged *ex vivo* culture, we reasoned that these functional and molecular abnormalities are likely to be cell-extrinsically mediated. Indeed, preLSC showed enrichment of gene-signatures associated with certain cell-extrinsic inflammatory mediators (TNFα, IFNγ, TGFβ, IL2) (Fig. 3k).

### Inflammation promotes *TP53-*associated clonal dominance

To understand the transcriptional signatures associated with leukaemic progression we analysed samples from 5 CP-MPN patients who subsequently developed *TP53-*sAML (“pre-*TP53*-sAML”) alongside 6 CP-MPN patients harbouring *TP53* mutated clones who remained in chronic phase (“CP *TP53-*MPN”, median 4.43 years [2.62-5.94] of follow-up, Fig.4a, Extended Data Fig.8). Compared to *TP53-*sAML samples, CP *TP53*-MPN had a lower VAF and number of *TP53* mutations (Extended Data Fig.8a-d). The type, distribution and pathogenicity score of *TP53* mutations were similar between chronic and acute stages (Extended Data Fig.8e,f). 5 pre-*TP53*-sAML samples and 4 CP *TP53*-MPN were analysed by TARGET-seq (Fig.4a). HSPC immunophenotype was similar for pre-*TP53*-sAML and CP *TP53-*MPN patients (Extended Data Fig.9a,b), and clearly distinct from *TP53*-sAML (Extended Data Fig.9b). Heterozygous *TP53* clones were identified in 3 pre-*TP53*-sAML patients and all 4 CP *TP53-*MPN (Fig.4b, Extended Data Fig.9c-k). A minor homozygous *TP53* mutated clone initially present in one CP *TP53*-MPN patient was undetectable after 4 years (Extended Data Fig.9f). As *TP53*-heterozygous mutant HSPCs represent the direct genetic ancestors of *TP53* “multi-hit” LSCs, we compared gene expression of heterozygous *TP53* mutant HSPC from pre-*TP53*-sAML (n=296) to CP *TP53*-MPN (n=314) (Fig.4b, blue boxes) to identify putative mediators of transformation. *TP53*-heterozygous HSPC from pre-*TP53*-sAML patients showed downregulation of TNFα and TGFβ associated gene signatures with upregulated expression of oxidative phosphorylation, DNA repair and interferon response genes (TableS5, Fig.4c, Extended Data Fig.9l), without changes in IFN receptor expression levels or concurrent interferon treatment (Extended Data Fig.9m, Table S1). This raises the possibility that inflammation might contribute to preleukaemic clonal evolution towards *TP53-*sAML.

**Fig. 4.**
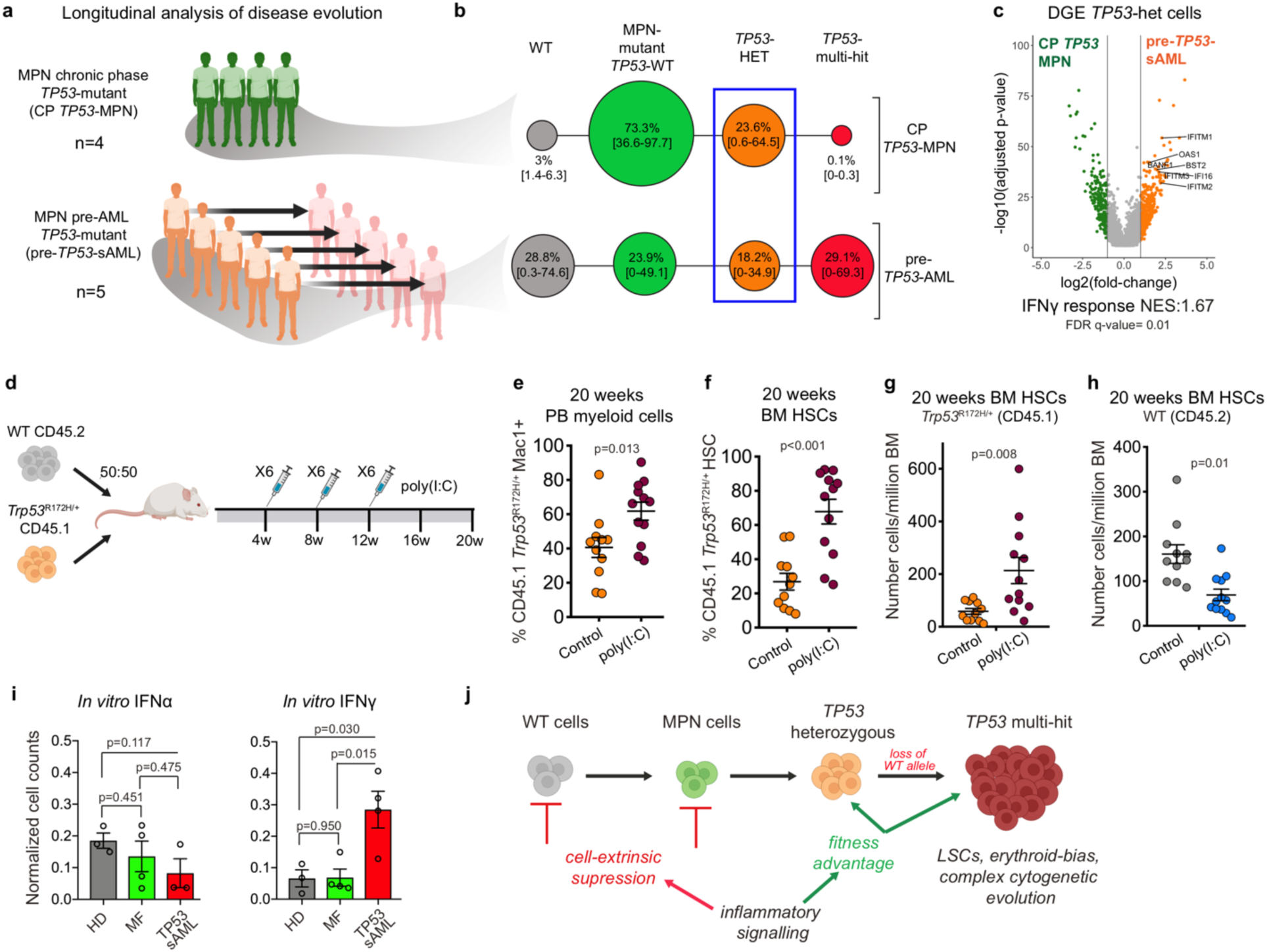
Inflammation promotes *TP53-*associated clonal dominance. **a,** Schematic study layout of the chronic phase and paired samples patient cohort selected for TARGET-seq analysis. **b,** Clonal evolution of *TP53*-mutant chronic phase patient samples without clinical evidence of transformation (CP *TP53*-MPN, n=4) and pre-*TP53*-sAML (patients who subsequently transformed to *TP53-*sAML) samples (n=5). The size of the circles is proportional to the average percentage of cells mapping to each clone, and each clone is coloured according to its genotype (related to Extended Data Fig.9c-k). *TP53*-heterozygous cells selected for subsequent transcriptional analysis are indicated by the blue box**. c,** Volcano plot of differentially expressed genes in *TP53*-mutant heterozygous cells from CP *TP53*-MPN (green; n=296 cells from 4 patients) and pre-*TP53*-sAML (orange; n=314 cells from 3 patients). Genes involved in the IFN-response are labelled, and GSEA analysis IFN-γ response Normalized Enrichment Score is indicated below. **d,** Experimental design of WT:*Trp53^R172H/+^* chimera serial poly(I:C) treatment. **e-h,** Analysis of chimera mice 20 weeks post-transplantation following 3 regimes of 6 poly(I:C) injections. Percentage of CD45.1 *Trp53^R172H/+^* Mac1^+^ cells in the peripheral blood **(e)** or bone marrow (BM) HSCs (Lin^-^Sca-1^+^c-Kit^+^CD150^+^CD48^-^) **(f)**, number of BM CD45.1 *Trp53^R172H/+^* HSC **(g)** and CD45.2 WT HSC **(h)** per million BM cells. n=11-12 mice per group in 2 independent experiments and 3 biological replicates. Mean ± s.e.m. is shown, and “p” indicates two-tailed unpaired t-test p-value. **i,** Percentage of viable cells after 12 days of liquid culture of Lin^-^ CD34^+^ cells from HD (n=3), MF (n=4) and *TP53*-sAML patients (n=3-4) treated with IFN-α (left) or IFN-γ (right). Number of cells were normalized to control condition. Barplot indicates mean ± s.e.m. from 5 independent experiments and “p”, two-tailed unpaired t-test p-value. **j,** Schematic proposed model of *TP53*- mutant driven transformation in MPN.

To evaluate the role of inflammation in *TP53-*driven leukaemia progression, we performed competitive mouse transplantation experiments between CD45.1+ *Trp53*^R172H/+^ and CD45.2+ *Trp53*^+/+^ BM cells followed by repeated poly(I:C) intraperitoneal injections, recapitulating chronic inflammation through induction of multiple pro-inflammatory cytokines^32, 33^ (Fig.4d). *Trp53* mutant peripheral blood myeloid cells, BM HSC (Lin^-^Sca1^+^c-Kit^+^CD150^+^CD48^-^) and LSK (Lin^-^Sca1^+^c-Kit^+^) were selectively enriched upon poly(I:C) treatment (Fig.4e,f, Extended Data Fig.10a-d). Crucially, the fitness advantage of *Trp53* mutant HSC and LSK was exerted both through an increase in numbers of *Trp53*^R172H/+^ HSPCs and reduction in numbers of wild-type competitors (Fig.4g,h, Extended Data Fig.10e,f). Treatment with poly(I:C) induced high levels of IFNγ (Extended Data Fig.10g), which is also increased in the serum of patients with MPN^34^. Treatment of human HSPC *in vitro* with IFNγ, but not IFNα, revealed selective resistance of *TP53-*sAML cells to IFNγ mediated proliferation inhibition compared to HD or MF cells, without changes in apoptosis (Fig.4i, Extended Data Fig.10h). Together, these results suggest that chronic inflammation favours the survival of *TP53* mutated cells whilst suppressing wild-type haematopoiesis, ultimately promoting clonal expansion of *TP53* mutant HSPCs (Fig.4j).

## Discussion

Here, we unravel multi-layered genetic, cellular and molecular intratumoural heterogeneity in *TP53* mutation driven disease transformation through single-cell multi-omic analysis. Allelic resolution genotyping of leukaemic HSPCs revealed a strong selective pressure for gain of *TP53* missense mutation, loss of the *TP53* wild-type allele and acquisition of complex CNAs, including cases with parallel genetic evolution during *TP53-*sAML LSC expansion. Despite the known dominant negative and/or gain of function effect of certain *TP53* mutations^28, 35^, loss of the *TP53* wild-type allele, a genetic event associated with a particularly dismal prognosis^2^, confers additional fitness advantage to *TP53-*sAML LSCs. As CNA were universally present in *TP53-*sAML with a very high clonal burden, it is not possible, even with high-resolution single-cell analyses, to disentangle the impact of *TP53-*multi hit mutation versus the effects of patient-specific CNA which were inextricably linked in all patients analysed.

Three distinct clusters of HSPCs were identified in *TP53-*sAML, including one characterized by overexpression of erythroid genes, of particular note as erythroleukaemia is a rare entity, associated with adverse outcome and *TP53* mutation^36, 37^. Analysis of a large AML cohort also revealed overexpression of erythroid genes as a more widespread phenomenon in *TP53* mutant AML, with disrupted balance of *GATA1* and *CEBPA* expression. Notably, *CEBPA* knockout or mutation is reported to cause a myeloid to erythroid lineage switch with increased expression of *GATA1*^29, 30^. Importantly, our data do not distinguish whether this lineage-switch is primarily an instructive versus permissive effect of *TP53-*mutation^38^. A second ‘*TP53-*sAML LSC’ cluster allowed us to establish a novel p53LSC-signature, which we demonstrated to be highly relevant to predict outcome in AML, independently of *TP53* status. This powerful approach could be more broadly applied in cancer, whereby single multi-omic cell derived gene scores can be used to stratify larger patient cohorts using bulk gene expression data.

A third *TP53* wild-type ‘preLSC’ HSPC cluster was characterized by quiescence signatures and defective differentiation, reflecting the impaired haematopoiesis observed in patients with *TP53-*sAML. Through integration of single cell multi-omic analysis with *in vitro* and *in vivo* functional assays we show that *TP53-*wild-type preLSCs are cell-extrinsically suppressed whilst chronic inflammation promotes the fitness advantage of *TP53* mutant cells, ultimately leading to clonal selection (Fig.4j). Inflammation is a cardinal regulator of HSC function with many effects on HSC fate and function^39^, including proliferation-induced DNA-damage and depletion of HSC^40^. There is emerging evidence that clonal HSC can become inflammation-adapted^39, 41, 42^ and by altering the response to inflammatory challenges, mutations can thus confer a fitness advantage to HSC. Here, we demonstrate a hitherto unrecognized effect of *TP53* mutations, which conferred a marked fitness advantage to HSPC in the presence of chronic inflammation, which we speculate could occur by altering the HSC response to proliferation-induced DNA-damage. Further studies are required to characterize this, and also the key inflammatory mediators involved, which we believe are unlikely to be restricted to a single axis, with a myriad of inflammatory mediators overexpressed in MPN^43^. Consequently, we believe that approaches which target the inflammatory state, rather than a specific cytokine, are likely to be required to restrain disease progression, as reported for bromodomain inhibitors^44^. Collectively, our findings provide a crucial conceptual advance relating to the interplay between genetic and non-genetic determinants of *TP53*-mutation associated disease transformation. This will facilitate the development of early detection and treatment strategies for *TP53-*mutant leukaemia. Since *TP53* is the most commonly mutated gene in human cancer^3, 45^, we anticipate these findings will be of broader relevance to other cancer types.

---

**Extended Data Fig.1.**
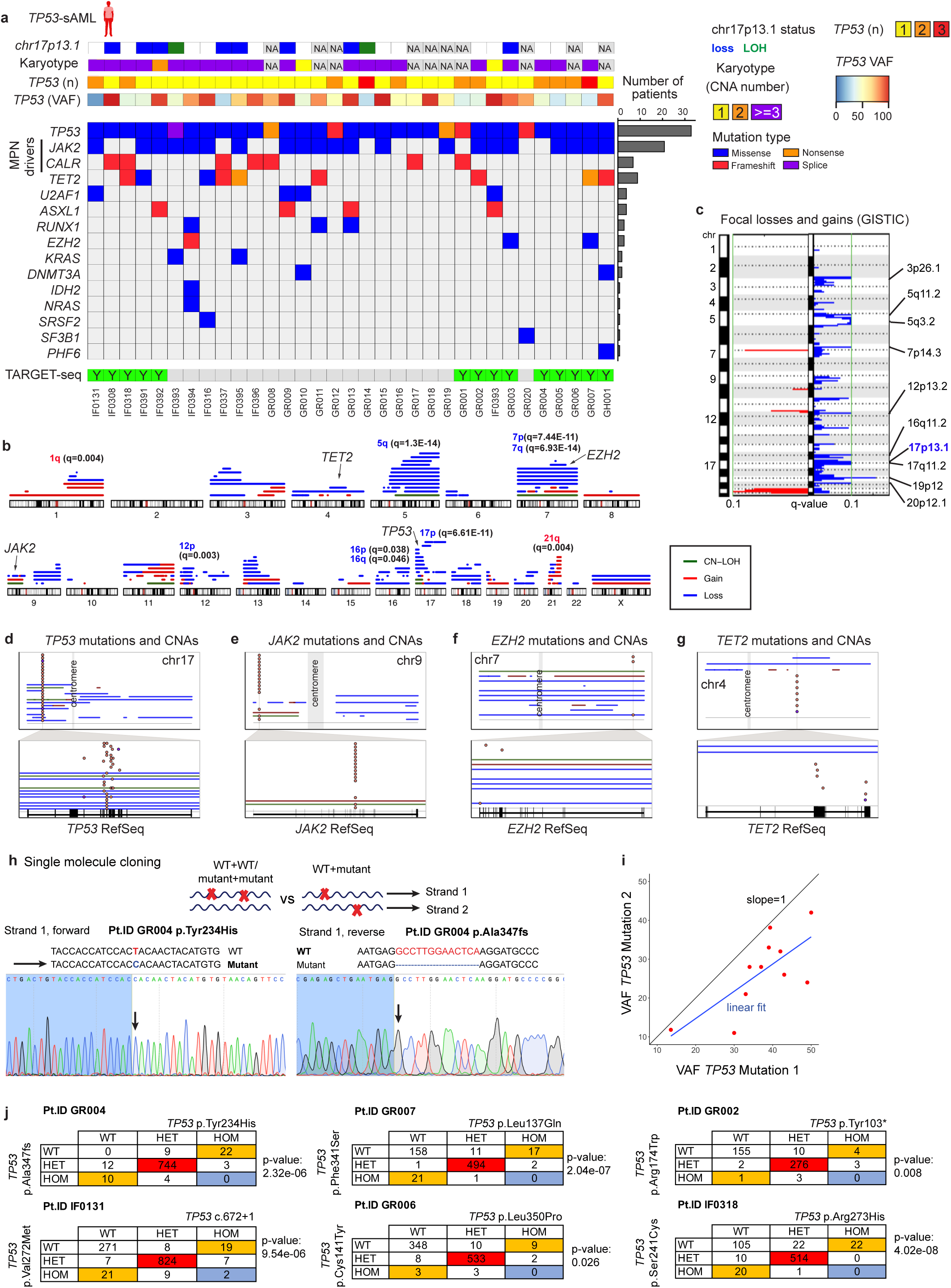
Genetic landscape of *TP53*-sAML. **a,** Mutations, CNAs, *TP53* VAF and allelic status identified in a cohort of 33 *TP53*-sAML patients by bulk sequencing. The barplot on the right indicates the frequency of each mutation in the cohort. The panel at the bottom indicates samples processed by TARGET-seq. **b-c,** Graphical representation of all CNAs identified by MoChA (**b**) and GISTIC analysis of recurrently lost (blue) and amplified (red) focal regions (**c**) in the same patients as in **(b)**. In **b**, GISTIC q-values of arm-level gains (red) and loses (blue) are indicated for each chromosome arm. In **c**, *TP53* chromosomal location is indicated in blue (17p13.1). **d-g,** Summary of CNA events spanning recurrently mutated genes *TP53* **(d)**, *JAK2* **(e)***, EZH2* **(f)** and *TET2* **(g)**, with evidence of deletion or loss of heterozygosity in the single-cell phylogenies computed in Extended Data Fig.2b-o. For each gene, top panel shows a whole chromosome view and the bottom one, the gene-level view and RefSeq track. Points indicate the location of each point mutation and solid lines indicate CNA status (blue:loss; red:gain; green:LOH). **h,** Sanger sequencing of single-molecule patient-derived *TP53* cDNA showing mutually exclusive alleles in the same cDNA molecule. **i,** VAF of *TP53* mutations in patients in which at least two *TP53* mutations were detected. Blue line represents the linear fit of the points, which deviates from the indicated slope that would be expected if mutations were on the same allele. When more than 2 mutations were present, the 2 with the highest VAF were analysed. **j,** Contingency table of *TP53* zygosity status in single cells from patients carrying two *TP53* mutations. Double-mutant heterozygous cells are coloured in red, mutually exclusive WT/homozygous or homozygous/WT genotypes in orange and homozygous/homozygous cells, in blue. “p” indicates exact binomial test p-value.

**Extended Data Fig.2.**
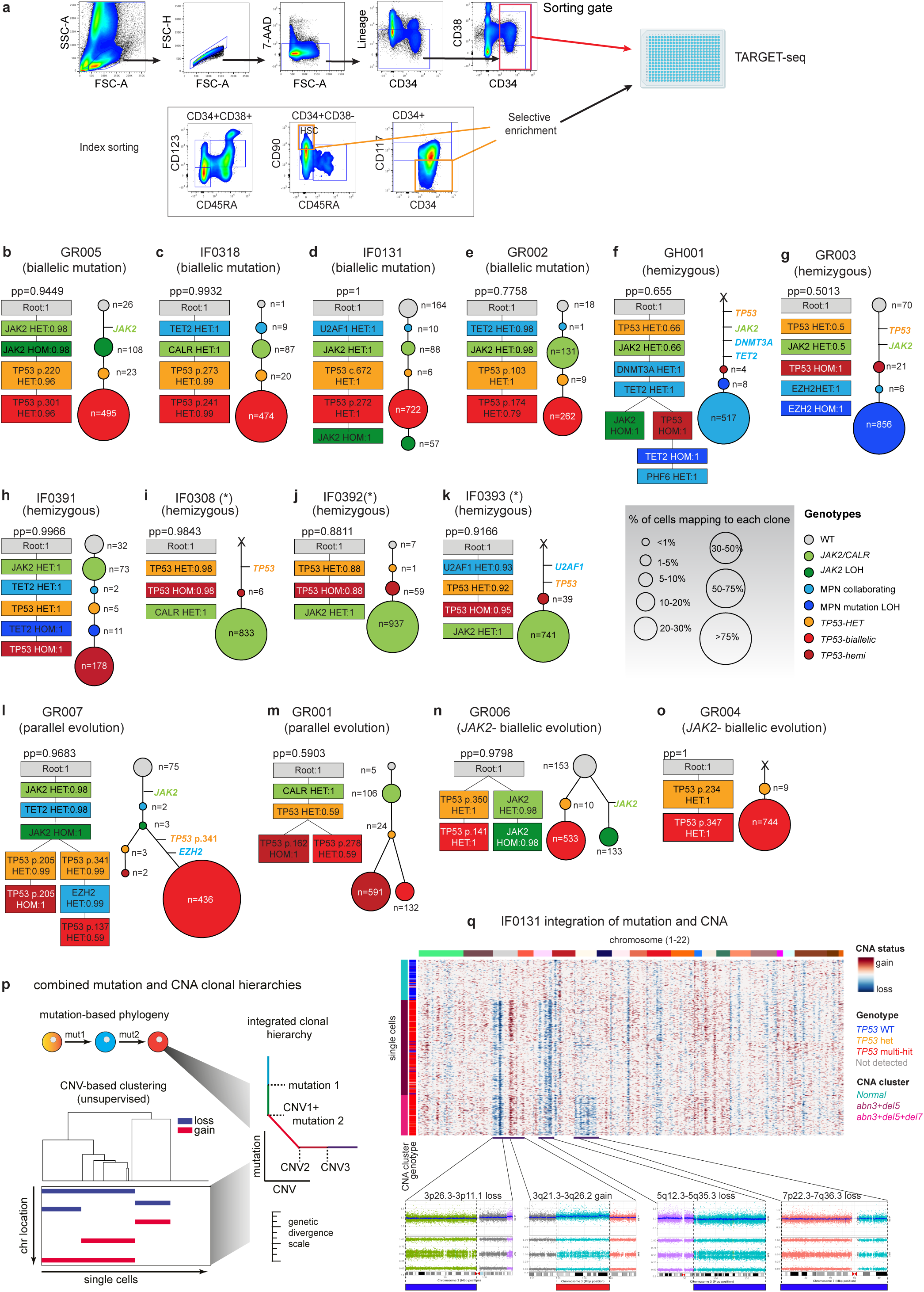
TARGET-seq sorting strategy and phylogenetic reconstruction of clonal hierarchies in *TP53-*sAML patients using a Bayesian model. **a**, Sorting strategy for TARGET-seq: Lineage^-^CD34^+^ cells were sorted into 384-well plates for subsequent library preparation. Selective enrichment of immunophenotypically defined populations (HSC: CD38^-^ CD90^+^CD45RA^-^; CD117^-^) is indicated with orange boxes. **b-o**, In each panel, corresponding to a different patient sample, the phylogenetic tree computed using SCITE is shown on the left and the number of cells mapping to each clone on the right. “pp” indicates the posterior probability of each consensus mutation tree, and the probability of each genotype transition is indicated inside each square for each mutation. The size of the circles is proportional to the size of each clone and is coloured according to the genotype indicated. The number of cells mapping to each clone is indicated in each circle and the type of *TP53* clonal evolution (biallelic mutation, hemizygous, parallel or *JAK2*-negative) below each patient’s ID. (*) indicates patients for which the high clonality of the sample prevented the faithful reconstruction of the order of mutation acquisition. Horizontal lines indicate mutation acquisition where none of the experimentally-detected clones matched that particular combination of mutations. Due to selective enrichment of certain subpopulations of cells (**a**), the numbers of cells assigned to each subclone in this figure is not necessarily representative of overall clonal burden, and early clones are likely over-represented due to selective enrichment of preleukemic HSCs. In contrast, the relative subclone percentages displayed in Fig.1 for the same patients have been corrected according to each populations’ frequency in the Lin^-^CD34^+^ compartment. **p**, Schematic representation of the strategy to reconstruct integrated clonal hierarchies based on single-cell TARGET-seq genotyping and inferCNV transcriptomic-based CNAs. **q**, Representative example of combined mutation and CNA hierarchies for patient IF0131, in which three cytogenetically-distinct subclones were detected. Corresponding cytogenetic lesions detected at the bulk level through high-density SNP arrays are shown in the bottom panels.

**Extended Data Fig.3.**
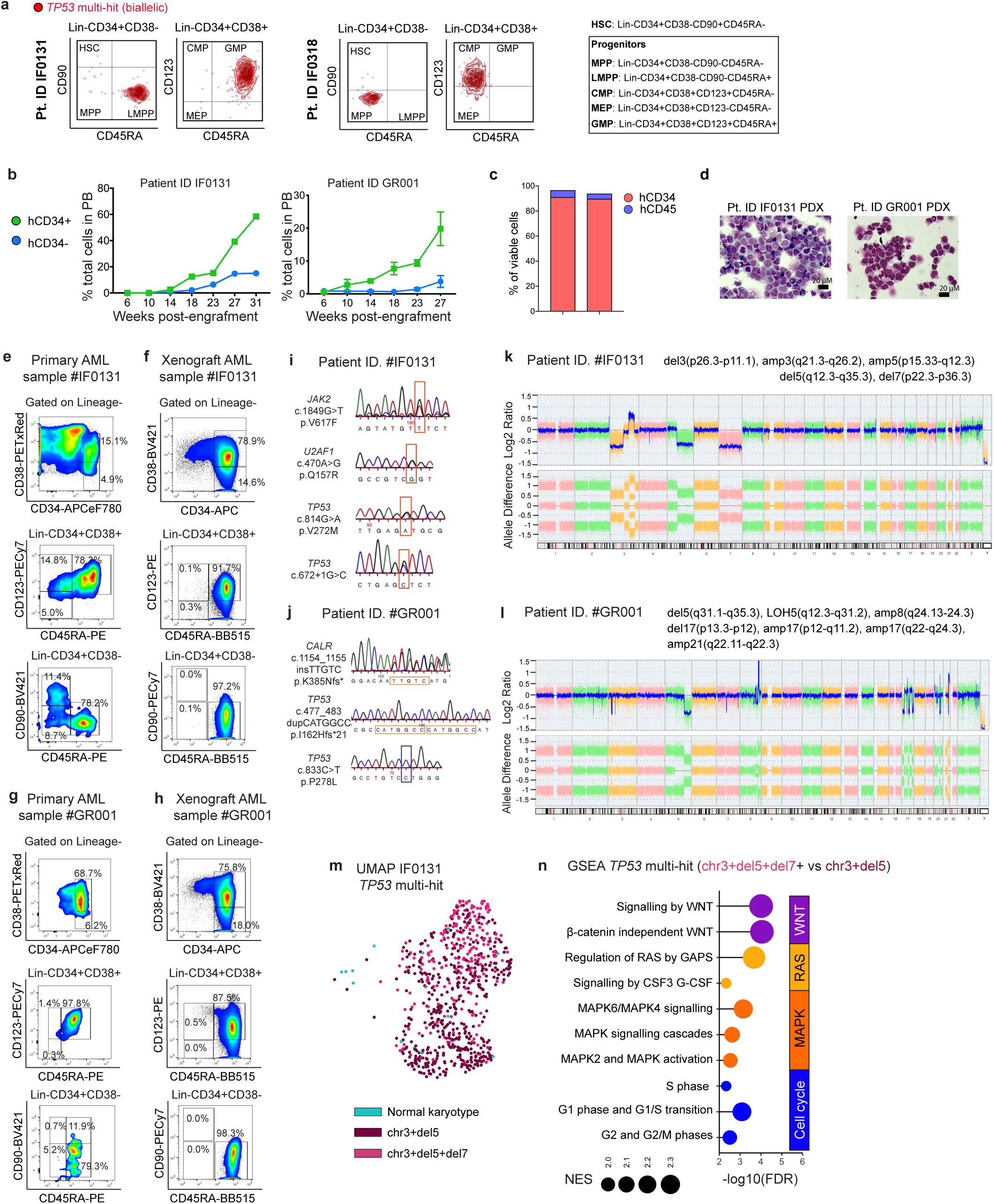
*TP53*-sAML xenograft characteristics. **a,** Integration of index sorting and single cell genotyping of *TP53* multi-hit HSPC from two representative patients. **b,** Serial readouts of human chimerism based on hCD34 and hCD45 expression in mouse peripheral blood (PB) for IF0131 (n=1) and GR001 (n=3, mean ± s.e.m. indicated). **c,** Proportion of hCD45 and hCD34- positive cells in total bone marrow (BM) from each PDX sample. **d,** Representative images from BM blasts isolated from PDX models **e-f,** Representative HSPC flow cytometry profiles of patient IF0131 PB mononuclear cells (MNCs) **(e)** and BM engrafted cells in immunodeficient mice at 31 weeks post transplantation **(f)**. **g-h,** Representative HSCP flow cytometry profiles of patient GR001 PB MNCs **(g)** and BM engrafted cells in immunodeficient mice at 27 weeks post-transplantation **(h)**. **i-l,** Mutations **(i,j)** and CNAs **(k,l)** detected in sorted LMPPs (Lin^-^CD34^+^CD38^-^ CD45RA^+^) from indicated PDX samples **(f,h)**. Boxes indicate location of each mutation (orange for mutant allele and blue, for wild-type) **m,** UMAP representation of *TP53* multi-hit cells from patient IF0131; cells are coloured according to their CNA status as in Fig.1g. **n,** GSEA analysis of cytogenetically distinct subclones in patient IF0131. Pathways enriched in *TP53* multi-hit abn3+del5+monosomy7 versus *TP53* multi-hit abn3+del5 Lin^-^CD34^+^ are shown and coloured according to pathway’s functional category. NES: Normalized Enrichment Score. FDR: False Discovery Rate.

**Extended Data Fig.4.**
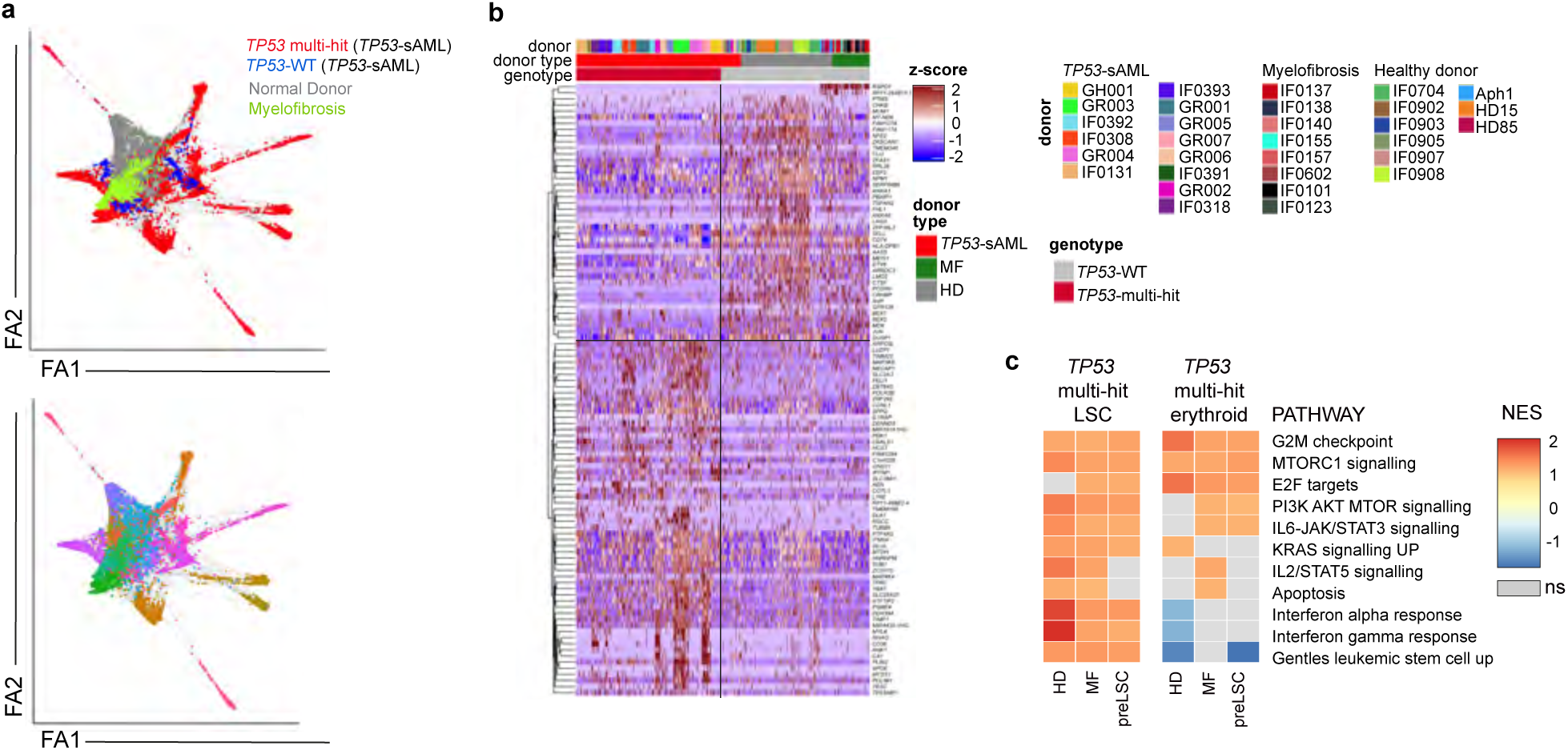
Single cell transcriptomic analysis of HD, MF and *TP53*-sAML HSPCs. **a,** Force-atlas representation of 17608 cells from HD (n=9), MF (n=8) and *TP53*-sAML patients (n=14; preleukemic: *TP53* WT, leukemic: *TP53* multi-hit) according to patient type (top) or donor (bottom). **b,** Heatmap of the top 100 differentially expressed genes identified between *TP53* multi-hit cells and preleukemic (*TP53*-WT; “preLSCs”), MF and HD cells. The type of donor, donor ID and *TP53* genotype is indicated on the top of the heatmap for each single cell. **c**, GSEA analysis of Lin-CD34+ *TP53* multi-hit LSC or erythroid-biased cells (Related to Fig.2a) from *TP53*-sAML patients, compared to Lin-CD34+ HD, MF and preLSCs. Heatmap indicates NES from selected genesets with FDR q- value<0.25. NES: Normalized Enrichment Score; FDR: False Discovery Rate. NS: non-significant.

**Extended Data Fig.5.**
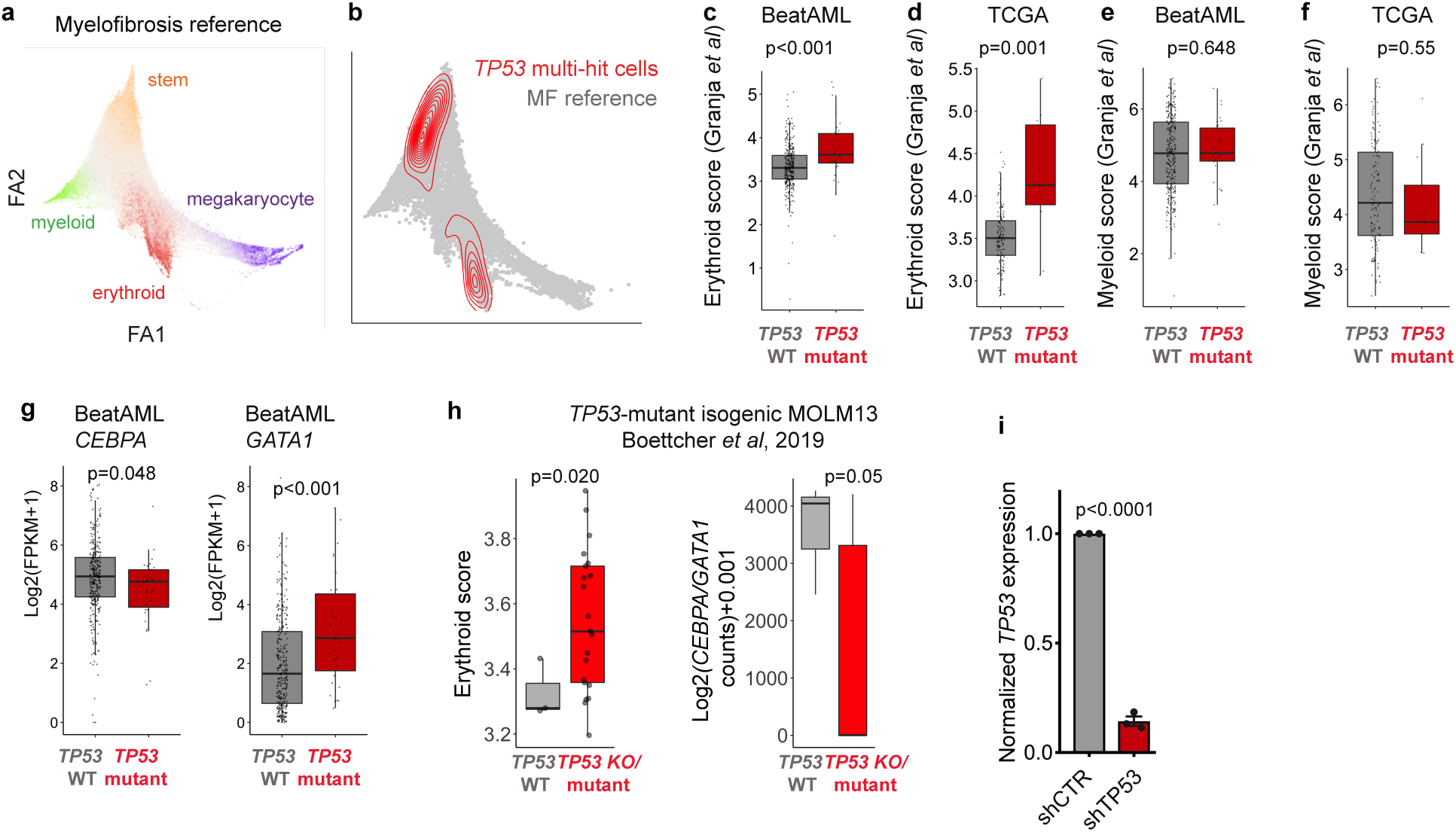
Aberrant erythroid differentiation in *TP53* mutant AML. **a-b,** Force Atlas representation of a CD34+ myelofibrosis (MF) atlas (**a**; Psaila et al, 2020) and latent-semantic index projection of *TP53* multi-hit cells from *TP53*-sAML patients into the MF cellular hierarchy (**b**). **c-g,** Expression of a second erythroid (**c,d**) and myeloid (**e,f**) gene score derived from a human hematopoietic atlas (Granja *et al*, 2019) and *GATA1/CEBPA* genes (**g**) in AML patients from BeatAML (**c,e,g**) and TCGA (**d,f**) datasets stratified by *TP53* mutational status (BeatAML: n=329 *TP53*-WT and n=31 *TP53*-mutant; TCGA: n=140 TP53-WT, n=11 *TP53*-mutant). Boxplots represent median and quartiles. “p” indicates Wilcoxon rank sum test p-values. **h,** Erythroid score (left) and *CEBPA/GATA1* gene expression ratios (right) in MOLM13 *TP53*-mutant isogenic cell lines (Boettcher *et al*, 2019). Boxplots represent median and quartiles. “p” indicates Wilcoxon rank sum test p-values. **i,** Fold-change *TP53* expression in CD34+GFP+ MPN primary cells following transduction with a lentiviral shRNA vector targeting *TP53* compared to a scramble control (shCTR). n=3 patients, 3 independent experiments. Barplot indicates mean ± s.e.m. and “p”, two-tailed unpaired t-test p-value.

**Extended Data Fig.6.**
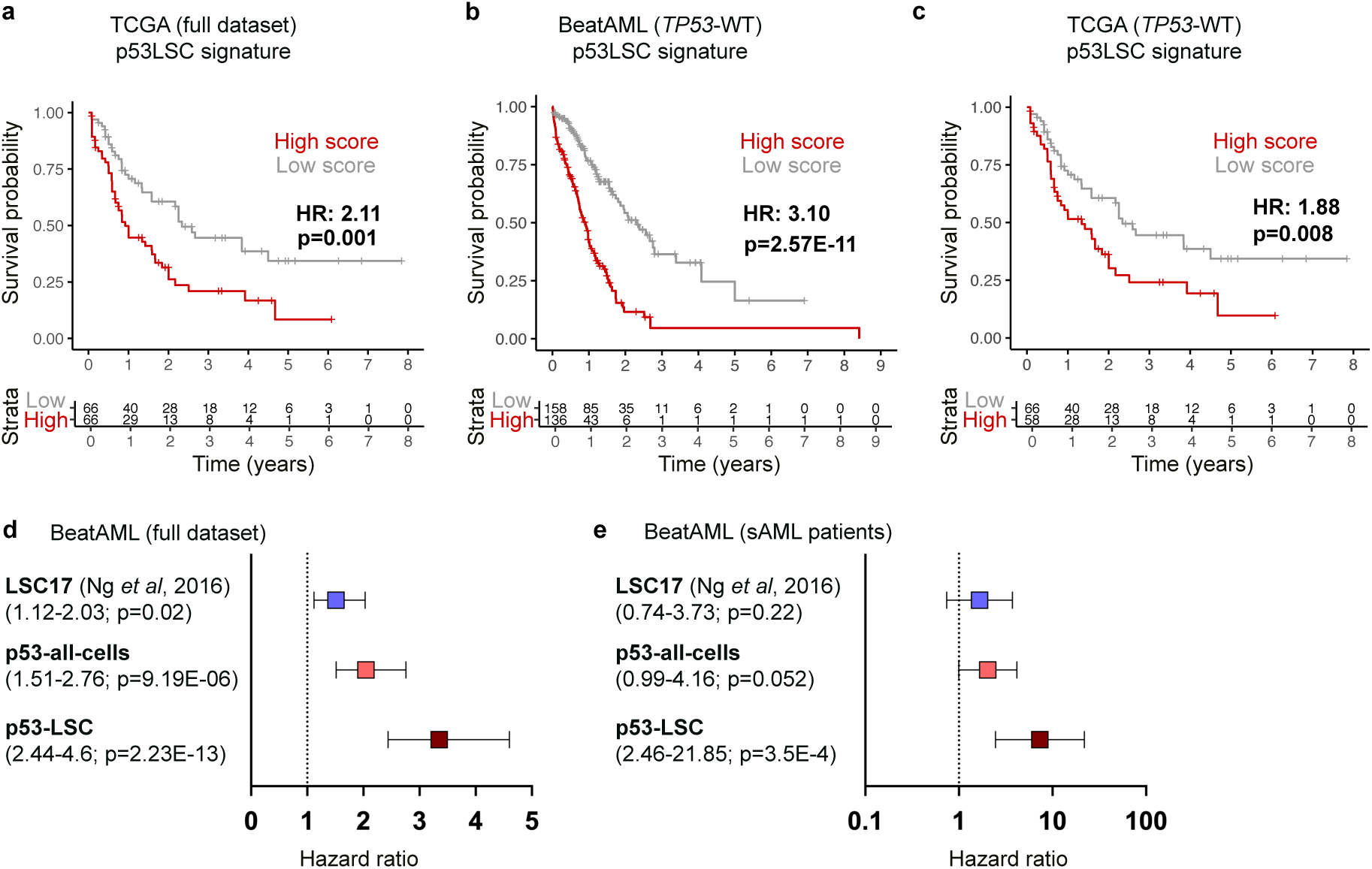
Validation of p53-LSC signature score in two independent cohorts. **a-c,** Kaplan-Meier analysis of *de novo* AML patients from the full TCGA AML dataset (n=132) (**a**), *TP53*-WT AML patients from BeatAML (n=294) (**b**) and *TP53*-WT AML patients from TCGA (n=124) (**c**) stratified according to high or low p53 LSC signature score. **d-e,** Hazard ratio (95% Cl) of all AML patients (n=322) (**d**) or secondary AML patients (n=49) (**e**) from the BeatAML cohort using LSC17 score (Ng et al, 2016), p53-all-cells score (derived from all *TP53*-mutant sAML cells) and p53-LSC signature score (derived from transcriptionally-defined LSCs; related to Fig.2a). Genes used for each score are listed in Table S4. “p” indicates log-rank test p-value.

**Extended Data Fig.7.**
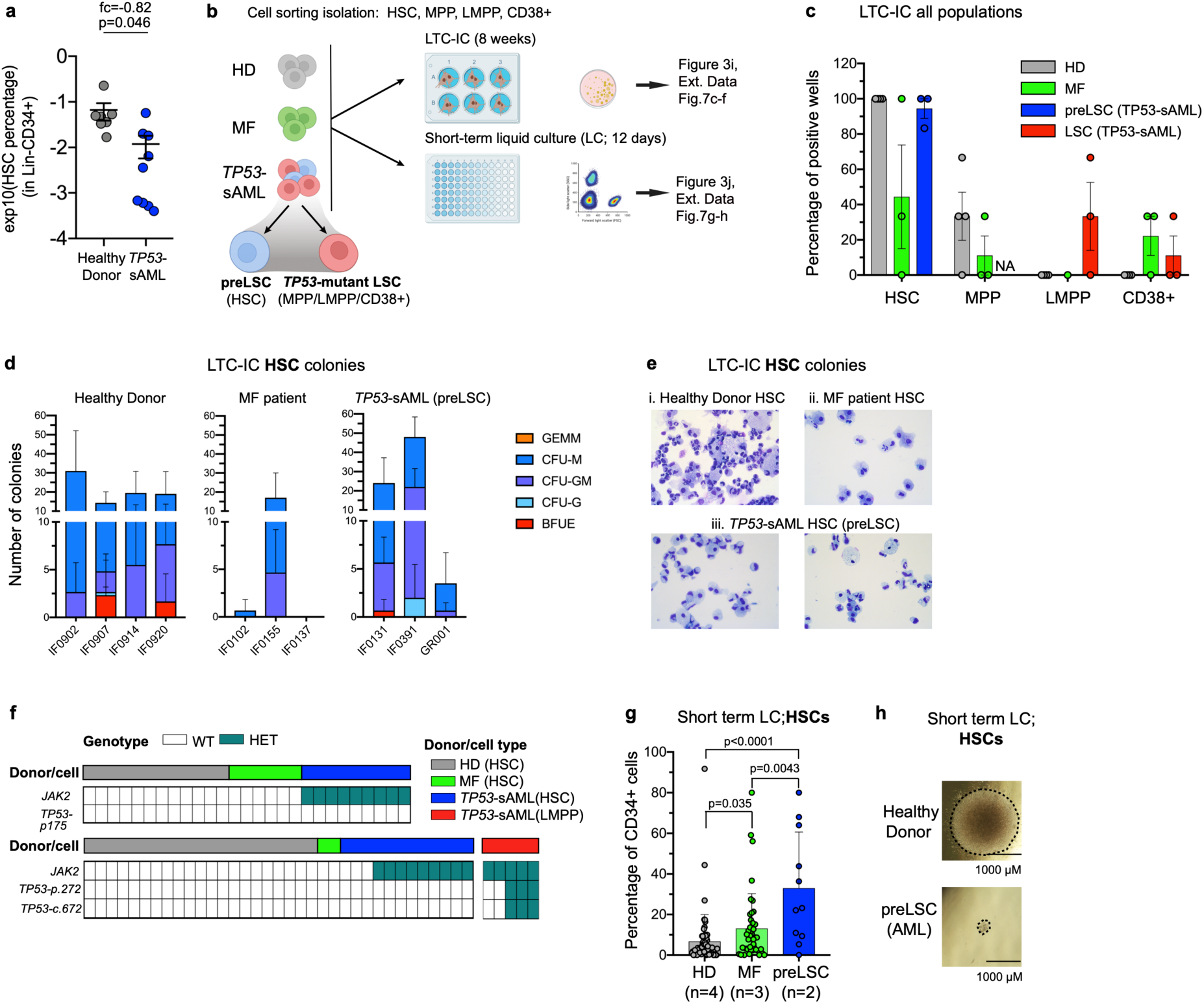
*In vitro* assessment of self-renewal and differentiation potential in preleukemic cells from TP53-sAML patients. **a**, Proportion of HSCs in mobilized PB or BM from healthy donors (n=7) and TP53-sAML patients (n=9) in which preLSCs were detected. Graph shows mean ± s.e.m, “p” indicates two-tailed Student t-test p-value and “fc”, fold-change. **b,** Schematic representation of Lin-CD34+ cell fractions isolated and *in vitro* assays performed. TP53-sAML patient samples used (n=3 : IF0131, IF0391, GR001) were known to have *TP53-WT* preleukemic stem cells (preLSC) in the HSC compartment (Related to Fig.3c). **c-d,** Long term culture-initiating cell *in vitro* assay. Percentage of positive wells in each immunophenotypic population (c) and clonogenic output **(d)** from HD (n=4), MF (n=3) and preLSCs from TP53-sAML (n=3). Barplot indicates mean ± s.e.m. from 2 independent experiments. e, Representative cytospin images of HSC-derived colonies from the same patient groups as in **(c-d). f,** Genotyping of HSC and LMPP-derived colonies from the same LTC-IC assay as in (c-e), demonstrating absence of *TP53* mutations in HSC-derived colonies, contrary to LMPPs. **g,** Percentage of CD34+ cells from HD (n=4), MF (n=3) and preLSCs from TP53-sAML (n=2) after 12 days of liquid culture in conditions promoting hematopoietic differentiation. Barplot indicates mean ± s.e.m from 3 independent experiments, and “p”, two-tailed Student t-test p-value. **h,** Representative image of liquid culture HSC-derived colonies for HD and TP53-sAML preLSCs, from the same experiment as in **(g).**

**Extended Data Fig.8.**
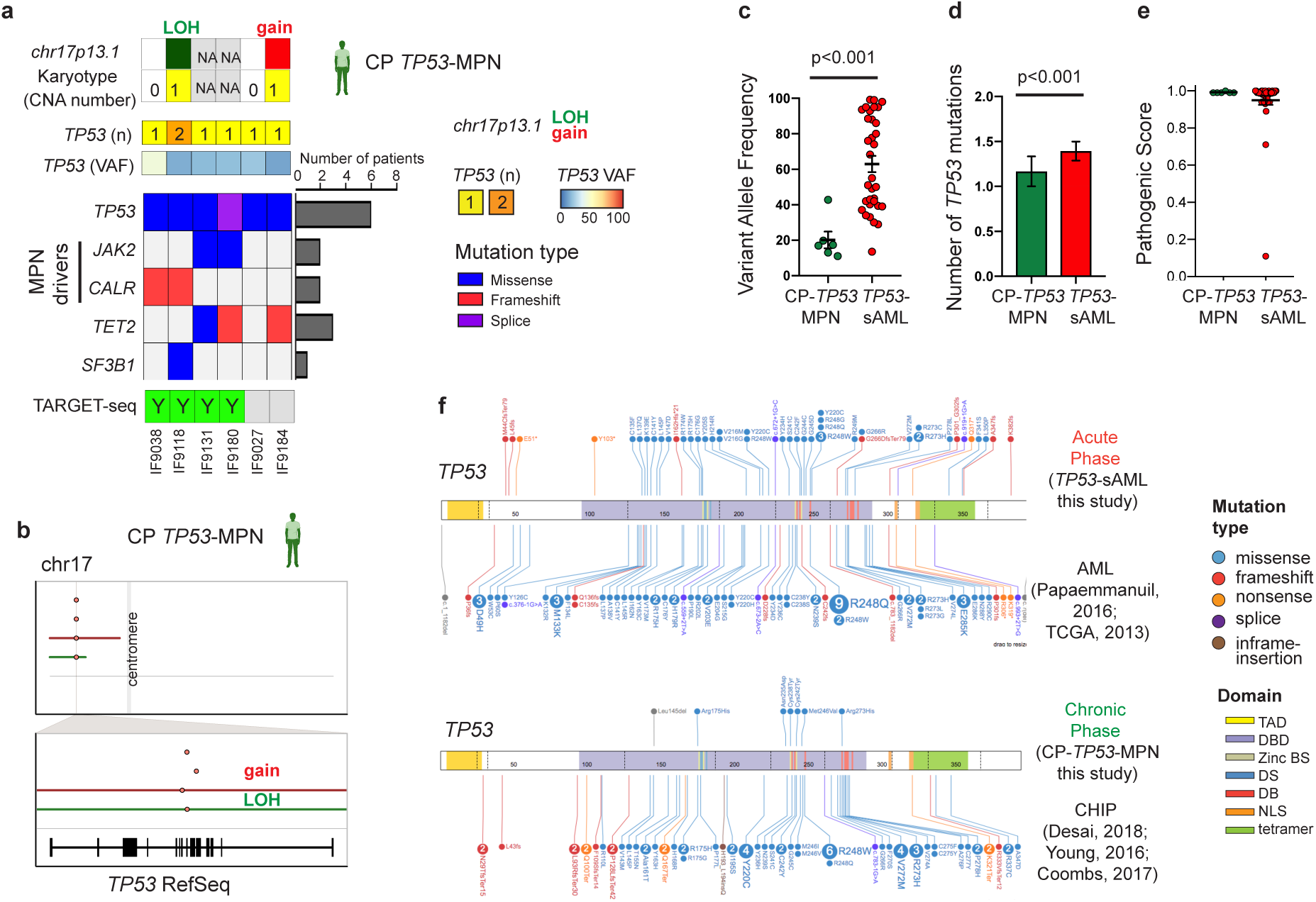
Genetic landscape of chronic phase *TP53*-mutant MPN. **a,** Point mutations and cytogenetic abnormalities identified in a cohort of 6 CP *TP53*-MPN patients with no evidence of clinical transformation after 4.43 years [2.62-5.94] median follow-up. The number of patients in which each gene is mutated in shown on the barplot on the right and patients processed for TARGET-seq analysis are indicated below the heatmap. **b,** Summary of CNA events in chr17 and *TP53* gene in the 2 CP *TP53*-MPN patients with detectable CNAs. The top panel shows a whole chromosome view and the bottom one, the gene-level view and RefSeq track. Points indicate the location of each point mutation and solid lines indicate CNA status. **c-e,** Comparison of variant allele frequency (**c**), number of TP53 mutations (**d**) and pathogenic scores (**e**) of *TP53* variants identified in CP *TP53*-MPN (n=6) and *TP53*-sAML patients (n=33). Mean ± s.e.m. is shown; “p” indicates two-tailed Mann-Whitney test p-value. **f,** Location and mutation type stratified by patient group (chronic/acute phase) as compared to previously published CHIP and AML patient cohorts.

**Extended Data Fig.9.**
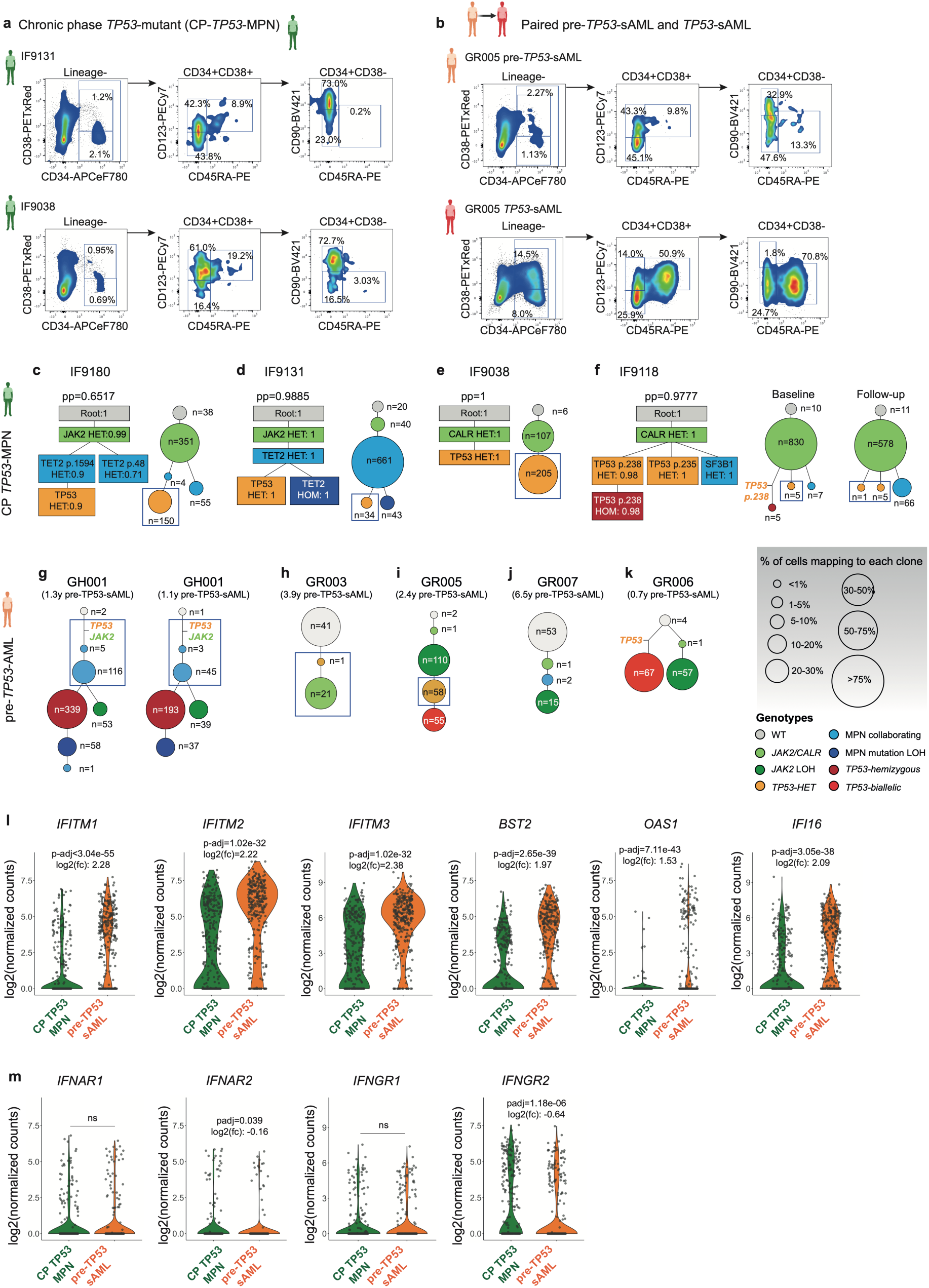
Clonal evolution and molecular signatures of *TP53*-mutant patients at chronic phase. **a-b,** Flow cytometry profiles of the Lin^-^CD34^+^ HSPC compartment in two CP *TP53*-MPN patients without evidence of clinical transformation (**a**) and in a representative paired chronic phase (**b**, up; pre-*TP53*-sAML) and acute phase (**b**, bottom; *TP53*-sAML) sample (Related to Fig.4a). **c-f,** Phylogenetic reconstruction of clonal hierarchies in CP *TP53*-MPN patients from single-cell TARGET-seq genotyping data. In each panel, the phylogenetic tree computed using SCITE is shown on the left, and the number of cells mapping to each clone for each patient, on the right. “pp” indicates posterior probability or each consensus mutation tree, and the probability of each genotype transition is indicated in the square for each mutation. The size of the circles is proportional to the size of each clone and is coloured according to the genotype indicated in the genotype key. For patient IF9118 **(f**), baseline (left) and 4 years of follow-up (right) samples are shown separately. **g-k,** Phylogenetic reconstruction of clonal hierarchies in pre-*TP53*-AML patients from single-cell TARGET-seq genotyping data (related to Extended Data Fig.2). The size of the circles is proportional to each clone’s size, and is coloured according to the genotype indicated in the genotype key. In panels (c-k), blue boxes indicate *TP53*-heterozygous clones used for the analysis presented in Fig.4c. **l-m,** Expression of key interferon-response genes **(l)** and interferon receptors **(m)** in *TP53*-heterozygous cells from CP *TP53*-MPN (n=296 cells) and pre-*TP53*-sAML patients (n=314 cells). “p-adj” indicates adjusted p-value from combined Fisher’s exact test and Wilcoxon tests, calculated using Fisher’s method and adjusted using Benjamini & Hochberg procedure; “fc” indicates fold-change (related to Fig.4c). Violin plots indicate log2(counts) distributions and each point represents the expression value of a single-cell.

**Extended Data Fig.10.**
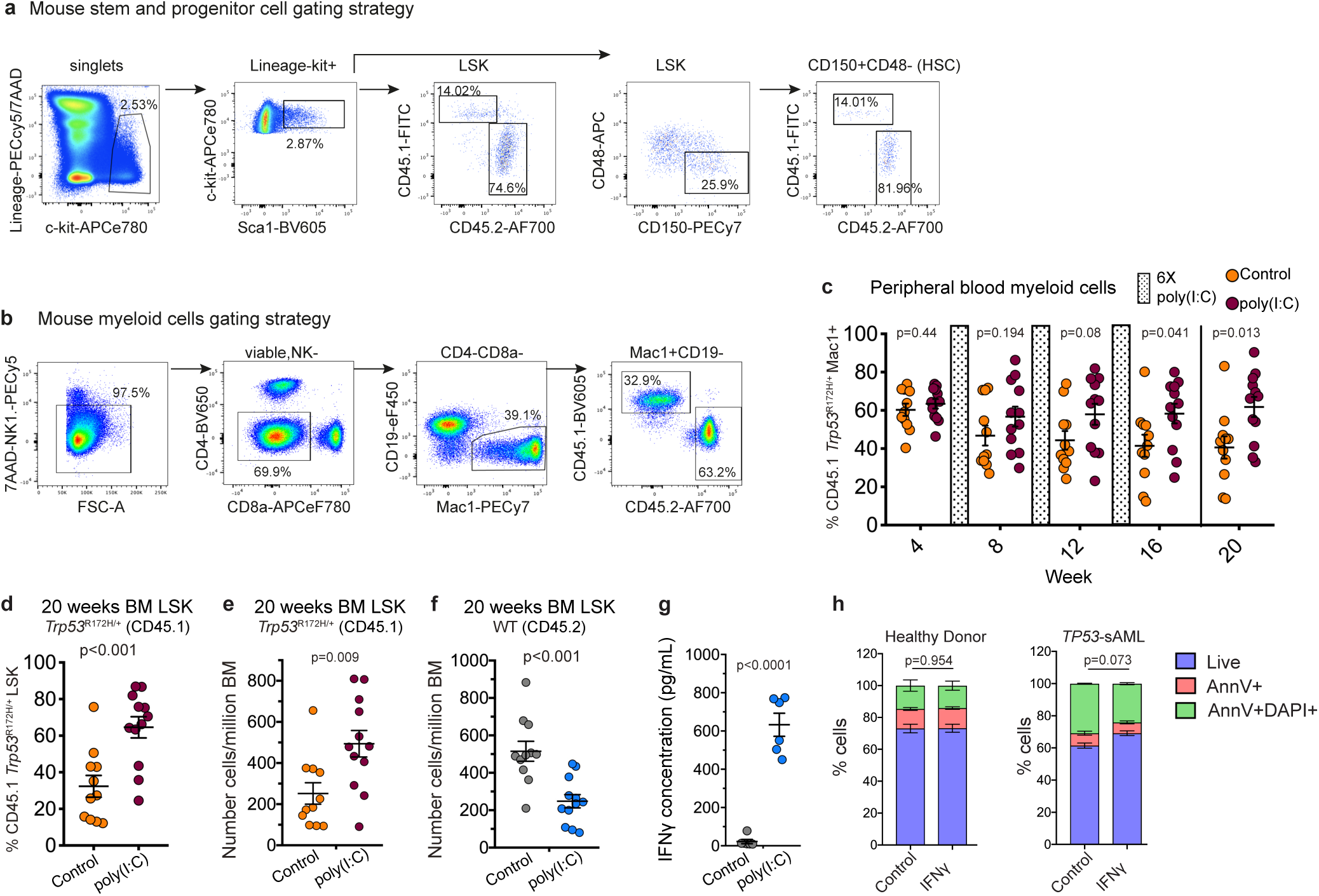
*TP53*-mutant cells display an aberrant inflammatory response. **a-b,** Gating strategy for mouse chimera experiments (Related to Fig.4d-h) used to quantify CD45.1+ LSK and HSCs populations in the BM (**a**) and myeloid cells in the peripheral blood (PB) (**b**). **c-f,** Analysis of WT:*Trp53^R172HI+^* chimera mice treated with 3 regimes of 6 poly(I:C) injections with serial readouts of CD45.1 *Trp53^R172HI+^* Mac1+ PB cells (**c**), percentage of CD45.1 *Trp53^R172HI+^* BM LSK (Lin-Sca-1+c-Kit+) (**d**), number of CD45.1 *Trp53^R172HI+^* BM LSK (**e**) and CD45.2 WT BM LSK per million BM cells (**f**) 20 weeks post transplantation. n=11-12 mice per group from 3 biological replicates in 2 independent experiments. Bars indicate mean ± s.e.m. and “p”, two-tailed unpaired t-test p-value. **g,** IFNy level in spleen serum 4h after poly(I:C) injection. n=6 mice per group from 2 independent experiments. Lines indicate mean ± s.e.m. and “p”, two-tailed unpaired t-test p-value. **h,** Percentage of apoptotic cells from healthy donor (n=3) or *TP53*-sAML patients (n=2) determined by Annexin-V/DAPI staining 24h after IFNy treatment of HSPCs. “p” indicates two-tailed unpaired t-test p-value.

## Methods

### Banking and processing of human samples

Primary human samples (peripheral blood or bone marrow, described in Table S1) were analysed with approvals from the Inserm Institutional Review Board Ethical Committee (project C19-73, agreement 21-794, CODECOH n°DC-2020-4324); and from the INForMeD Study (REC: 199833, 26 July 2016, University of Oxford). Patients and normal donors provided written informed consent in accordance with the Declaration of Helsinki for sample collection and use in research. For secondary AML patients, we specifically selected samples from patients with known *TP53*-mutation.

Cells were subjected to Ficoll gradient centrifugation and for some samples, CD34 enrichment was performed using immunomagnetic beads (Miltenyi). Total mononuclear cells (MNCs) or CD34^+^ cells were frozen in FBS supplemented with 10% DMSO for further analysis.

### Targeted bulk sequencing

Bulk genomic DNA from patient samples’ mononuclear or CD34^+^ cells was isolated using DNeasy Blood & Tissue Kit (Qiagen) or QIAamp DNA Mini Kit (Qiagen) as per manufacturer’s instructions. Targeted sequencing was performed using a TruSeq Custom Amplicon panel (Illumina) or a Haloplex Target Enrichment System (Agilent technologies) with amplicons designed around 32, 44 or 77 genes^46^. Targets were chosen based on the genes/exons most frequently mutated and/or likely to alter clinical practice (diagnostic, prognostic, predictive or monitoring capacity) across a range of myeloid malignancies (e.g. MDS/AML/MPN). Targets covered in all panels include *ASXL1, CALR, CBL, CEBPA, CSF3R DNMT3A, EZH2, FLT3, HRAS, IDH1, IDH2, JAK2, KIT, KRAS, MPL, NPM1, NRAS, PHF6, RUNX1, SETBP1, SF3B1, SRSF2, TET2, TP53, U2AF1, WT1, ZRSR2*. Sequencing was performed with a MiSeq sequencer (Illumina), according to the manufacturer’s protocols. Results were analysed after alignment of the reads using two dedicated pipelines, SOPHiA DDM^®^ (Sophia Genetics) and an in-house software GRIO-Dx^®^. For all samples, an average depth exceeding 200X for > 90% of the target regions was required, or as previously described^16^. All pathogenic variants were manually checked using Integrative Genomics Viewer software. Analysis is presented in Extended Data Fig.1a and Extended Data Fig.8a.

Pathogenic scores for each *TP53* variant (Extended Data Fig.8e) were derived from COSMIC (Catalogue Of Somatic Mutations In Cancer) using the FATHMM-MKL algorithm. The FATHMM-MKL algorithm integrates functional annotations from ENCODE with nucleotide-based hidden Markov models to predict whether a somatic mutation is likely to have functional, molecular and phenotypic consequences. Scores greater than 0.7 indicate that a somatic mutation is likely pathogenic, whilst scores less than 0.5 indicate a neutral classification.

The type and location of *TP53* mutations from this study, *de novo* AML patients and CHIP individuals represented in Extended Data Fig.8f were generated using Pecan Portal^47^. *De novo* AML *TP53* mutations were downloaded from Papaemmanuil, *et al.*^48^ and Ley, *et al.*^27^; CHIP associated *TP53* mutations were obtained from Coombs, *et al.,* Desai, *et al.*, Young, *et al.* ^49–51^

### Sanger sequencing of patient-associated mutations in PDX models

Genomic DNA from PDX sorted populations (LMPP: hCD45^+^Lin^-^CD34^+^CD38^-^ CD45RA^+^CD90^-^ and GMP: hCD45^+^Lin^-^CD34^+^CD38^+^CD45RA^+^CD123^+^) was extracted using QIAamp DNA Mini Kit (Qiagen). Sanger sequencing was performed with forward or reverse primers (TableS6a) targeting mutations identified by targeted bulk sequencing in the corresponding primary samples using Mix2seq kit (Eurofins Genomics) and sequences were analysed with the ApE editor.

### Single Nucleotide Polymorphism Array sample preparation, Copy Number Variant and Loss of Heterozygosity Analysis

Bulk genomic DNA from patients’ mononuclear cells was isolated using DNeasy Blood & Tissue Kit (Qiagen) as per manufacturer’s instructions. 250 ng of gDNA were used for hybridization on an Illumina Infinium OmniExpress v1.3 BeadChips platform.

To call mosaic copy number events in primary patient samples, genotyping intensity data generated was analysed using the Illumina Infinium OmniExpress v1.3 BeadChips platform. Haplotype phasing, calculation of log R ratio (LRR) and B-allele frequency (BAF) and calling of mosaic events was performed using Mocha (Mocha: A BCFtools extension to call mosaic chromosomal alterations starting from phased VCF files with either B Allele Frequency (BAF) and Log R Ratio (LRR) or allelic depth (AD)), as previously described^52, 53^. In brief, Mocha comprises the following steps: (1) filtering of constitutional duplications; (2) use of a parameterized hidden Markov model to evaluate the phased BAF for variants on a per-chromosome basis; (3) deploying a likelihood ratio test to call events; (4) defining event boundaries; (5) calling copy number; (6) estimating the cell fraction of mosaic events. A series of stringent filtering steps was applied to reduce the rate of false positive calls. To eliminate possible constitutional and germline duplications, excluding calls with lod_baf_phase <10, those with length <500KBP and rel_cov>2.5, and any gains with estimated cell fraction >80%, logR>0.5 or length <24Mb. Given that interstitial LOH are rare and likely artefactual, all LOH events <8Mb were filtered^52^. Events on genomic regions reported to be prone to recurrent artefact^52^ (chr6<58Mb, chr7>61Mb, and chr2 >50Mb) were also filtered, and those where manual inspection demonstrated noise or sparsity in the array.

To find common genomic lesions on a focal and arm level, Infinium OmniExpress arrays were initially processed with Illumina Genome Studio v2.0.4. Following this, Log R Ratio (LRR) data was extracted for all probes and array annotation obtained from Illumina (InfiniumOmniExpress-24v1-3_A1). LRR data was then smoothed and segmentation called using the CBS algorithm from the DNACopy^54, 55^ v1.60.0 package in R. A minimum number of 5 probes was required to call a segment, and segments where analysed using GenomicRanges^56^ v1.38.0. Definitions of amplification, gain, loss and deletion events where as outlined in Bashton, *et al.*^57^. Segmentation data was then analysed in GISTIC^58^ v2.023.

For PDX models, genomic DNA from sorted populations (LMPP: hCD45^+^Lin^-^CD34^+^CD38^-^ CD45RA^+^CD90^-^ and GMP: hCD45^+^Lin^-^CD34^+^CD38^+^CD45RA^+^CD123^+^) was extracted using QIAamp DNA Mini Kit (Qiagen). SNP-CGH array hybridization was performed using the Affymetrix Cytoscan® HD (Thermo Fisher Scientific) according to the manufacturer’s recommendations. DNA amplification was checked using BioSpec-nano^TM^ spectrophotometer (Shimadzu) with expected concentrations between 2,500 and 3,400ng/μL. DNA length distribution post-fragmentation was checked using D1000 ScreenTapes on Tapestation 4200 instrument (Agilent Technologies). Cytoscan HD array includes 2.6 million markers combining SNP and non-polymorphic probes for copy number evaluation. Raw data CEL files were analysed using the Chromosome Analysis Suite software package (v4.1, Affymetrix) with genome version GRCh37 (hg19) only if achieving the manufacturer’s quality cut-offs. Only CNAs > 10kb were reported in the analysis presented in Extended Data Fig.3k,l.

### Single-molecule cloning and sequencing of patient-derived cDNA

To experimentally verify the biallelic nature of *TP53* mutations in *TP53*-sAML patients, cDNA from a selected patient with putative *TP53* biallelic status (Patient ID GR004) was PCR-amplified using cDNA-specific primers spanning both *TP53* mutations (Fwd: 5’- GACCCTTTTTGGACTTCAGGTG-3’, Rev: 5’-CCATGAGCGCTGCTCAGATAG-3’). PCR amplification was performed with KAPA 2X Ready Mix (Roche), a Taq-derived enzyme with A-tailing activity, for direct cloning into a TA vector (pCR2.1 TOPO vector, TOPO® TA Cloning® Kit, Invitrogen) as per manufacturer’s instructions. Sanger sequencing for 10 different colonies was performed using M13 forward and reverse primers; a representative example is shown in Extended Data Fig.1h.

### Fluorescent activated cell sorting (FACS) and single-cell isolation

Single cell FACS-sorting was performed as previously described^16^, using BD Fusion I and BD Fusion II instruments (Becton Dickinson) for 96-well plate experiments or bulk sorting experiments, and SH800S or MA900 (SONY) for 384-well plate experiments. Experiments involving isolation of human haematopoietic stem and progenitor cells (HSPCs) included single colour stained controls (CompBeads, BD Biosciences) and Fluorescence Minus One controls (FMOs). Antibodies used for HSPC staining are detailed in TableS7a (Panel A or B).

Briefly, single cells directly sorted into 384-well plates containing 2.07 μL of TARGET-seq lysis buffer^59^. Lineage^-^CD34^+^ cells were indexed for CD38, CD90, CD45RA, CD123 and CD117 markers, which allowed us to record the fluorescence levels of each marker for each single cell. 7- aminoactinomycin D (7-AAD) was used for dead cell exclusion. Flow cytometry profiles of the HSPC compartment (Extended Data Fig.2, Fig.9) were analysed using FlowJo software (version 10.1, BD Biosciences).

### Single-cell TARGET-seq cDNA synthesis

RT and PCR steps were performed as previously described^59^, using 24 cycles of PCR amplification. Target-specific primers spanning patient-specific mutations were added to RT and PCR steps (TableS6a). After cDNA synthesis, cDNA from up to 384 single-cell libraries was pooled, purified using Ampure XP Beads (0.6:1 beads to cDNA ratio; Beckman Coulter) and resuspended in a final volume of 50 μL of EB buffer (Qiagen). The quality of cDNA traces was checked using a High Sensitivity DNA Kit in a Bioanalyzer instrument (Agilent Technologies).

### Whole transcriptome library preparation and sequencing

Pooled and bead-purified cDNA libraries were diluted to 0.2 ng/μL and used for tagmentation-based library preparation using a custom P5 primer and 14 cycles of PCR amplification^59^. Each indexed library was purified twice with Ampure XP beads (0.7:1 beads to cDNA ratio), quantified using Qubit dsDNA HS Assay Kit (Invitrogen, Cat# Q32854) and diluted to 4 nM. Libraries were sequenced on a HiSeq4000, HiSeqX or NextSeq instrument using a custom sequencing primer for read1 (P5_seq: GCCTGTCCGCGGAAGCAGT GGTATCAACGCAGAGTTGC*T, PAGE purified) with the following sequencing configuration: 15 bp R1; 8 bp index read; 69 bp R2 (NextSeq) or 150 bp R1; 8 bp index read; 150 bp R2 (HiSeq).

### TARGET-seq single-cell genotyping

After RT-PCR, cDNA+amplicon mix was diluted 1:2 by adding 6.25 μL of DNAse/RNAse free water to each well of each 384-well plate. Subsequently, a 1.5 μL aliquot from each single cell derived library was used as input to generate a targeted and Illumina-compatible library for single cell genotyping^59^. In the first PCR step, target-specific primers containing a plate-specific barcode (TableS6b) were used to amplify the target regions of interest. In a subsequent PCR step, Illumina compatible adaptors (PE1/PE2) containing single-direction indexes (Access Array™ Barcode Library for Illumina® Sequencers-384, Single Direction, Fluidigm) were attached to pre-amplified amplicons, generating single-cell barcoded libraries. Amplicons from up to 3,072 libraries were pooled and purified with Ampure XP beads (0.8:1 ratio beads to product; Beckman Coulter). These steps were performed using Biomek FxP (Beckman Coulter), Mosquito (TTP Labtech) and VIAFLO 96/384 (INTEGRA Biosciences) liquid handling platforms. Purified pools were quantified using Qubit dsDNA HS Assay Kit (Invitrogen, Cat# Q32854) and diluted to a final concentration of 4 nM. Libraries were sequenced on a MiSeq or NextSeq instrument using custom sequencing primers as previously described^59^ with the following sequencing configuration: 150 bp R1; 10 bp index read; 150 bp R2.

### Targeted single-cell genotyping analysis

#### Data pre-processing

For each cell, the FASTQ file containing both targeted gDNA and cDNA-derived sequencing reads was aligned to the human reference genome (GRCh37/hg19) using Burrow-Wheeler Aligner (BWA v0.7.17)^31^ and STAR (v2.6.1d)^60^. Custom perl scripts were used to demultiplex the gDNA and mRNA reads in the BAM file into separate SAM files based on targeted-sequencing primer coordinates (https://github.com/albarmeira/TARGET-seq). Next, Samtools (v1.9)^61^ was used to concatenate the BAM header to the resulting SAM files before re-converting the SAM file to BAM format, which was subsequently sorted by genomic coordinates and indexed. Both gDNA and mRNA reads were tagged with the cell’s unique identifier using Picard (v2.3.0) “*AddOrReplaceReadGroups”* and duplicate reads were subsequently marked using Picard “*MarkDuplicates”*. The sequencing reads overhanging into intronic regions in the mRNA reads were additionally hard-clipped using GATK (v4.1.2.0) *SplitNCigarReads*^62, 63^.

#### Variant calling

Variants were called from the processed BAM files using GATK *Mutect2* with the options [*--tumor-lod-to-emit 2.0 --disable-read-filter NotDuplicateReadFilter --max-reads-per-alignment-start*] to increase the sensitivity of detecting low-frequency variants. The frequency of each nucleotide (A, C, G, T) and indels at each pre-defined variant site were also called using a Samtools *mpileup* as previously described^16^. Lastly, the coverage at each pre-defined variant site were computed using Bedtools (v2.27.1)^64^.

To determine the coverage threshold of detection for each variant site, the coverage for “blank” controls (empty wells) were first tabulated. A cut-off coverage outlier value was computed as having a coverage exceeding 1.5 times the length of the interquartile range from the 75th percentile. Next, a value of 30 was added to this outlier value to yield the final coverage threshold to be used for genotype assignment.

#### Genotype assignment

For each pre-defined variant site, the number of reads representing the reference and alternative (variant) alleles for indels (insertion and deletions) and SNVs (single nucleotide variants) were tabulated from the outputs of GATK *Mutect2* and Samtools *mpileup*, respectively.

Here, a genotype scoring system was introduced to assign each variant site into one of three possible genotypes: wildtype, heterozygous, or homozygous mutant. Chi-square (χ^!^) test was first used to compare the observed frequency of reference and alternative alleles against the expected fraction of reference and alternative alleles corresponding to the three genotypes. The expected fraction of the reference alleles was 0.999, 0.5, and 0.001, and the expected fraction of the alternative alleles was 0.001, 0.5, and 0.999 for wildtype, heterozygous, and homozygous mutant genotype, respectively. The χ^!^ Statistics were then tabulated for each fitted model and converted to genotype scores using the following formula:

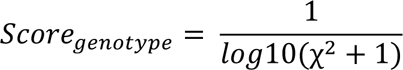

The genotype assigned to the variant site was based on the genotype model with the highest score.

Next, the variant (alternative) allele frequency (VAF) was computed and variant sites with 2 < VAF < 4 and 96 < VAF < 98 were reassigned as “ambiguous”. For cells with no variants detected at the specific variant sites by the mutation callers (either due to the absence of the variants, i.e. wild-type genotype, or that such variants were present below the detection limit), a “wild-type” genotype was assigned to those cells with a coverage above the specific threshold and “low coverage” to those cells with coverage below such threshold.

Taken together, each variant site was assigned one of the five following genotypes: wildtype, heterozygous, homozygous mutant, ambiguous, or low coverage. Variants with ambiguous or low coverage assignments for a particular cell were excluded from the analysis.

### Computational reconstruction of clonal hierarchies

Genotypes for each single cell were recoded for input to SCITE in a manner inspired by Morita *et al* ^65^: each mutation in each gene was coded as two loci, representing two different alleles. In the first recorded loci, all homozygous calls from each mutation where coded as heterozygous genotype calls. In the second recorded loci, all heterozygous and homozygous genotype calls in the original mutation matrix were coded as homozygous reference (i.e. WT) and heterozygous, respectively. For example, if for a certain mutation 0 represents WT status, 1 encodes heterozygous and 2 refers to homozygous status, these would be encoded as (0,0), (1,0) and (1,1) respectively, where the first term in the parenthesis corresponds to the first loci and the subsequent, to the second loci.

Then, SCITE was used (git revision 2016b31, downloaded from https://github.com/cbg-ethz/SCITE.git66) to sample 1000 mutation trees from the posterior for every single-cell genotype matrix corresponding to a particular patient, where all possible mutation trees are equally likely *a priori*. For patients in which several disease timepoints were available, all timepoints were merged for SCITE analysis. As parameters for every SCITE run “–fd 0.01” (corresponding to the allelic dropout rate of reference allele in our adapted SCITE model), “-ad 0.01” (corresponding to the allelic dropout of the alternate allele), a chain length (-l) of 1e6, and a thinning interval of 1 while marginalizing out cell attachments (-p 1 -s) were used.

To summarize the posterior tree sample distribution, the number of times a particular sample matched each tree was computed. For each patient, the most common tree topology in the posterior tree samples is reported (Extended Data Fig.2b-o, Fig.9c-k), where “pp” is the proportion of samples that match this tree. For each clade in the most common posterior tree, clade probabilities were estimated as the proportion of trees in the posterior that contained the clade. These are indicated in each square for each mutation in (Extended Data Fig.2b-o, Fig.9c-k).

#### Clone assignment

For every patient’s most common posterior tree, we assigned every cell to the tree node that matches the genotype of that particular cell. If an exact match was not found, then for every tree node the loss of assigning a cell to that node was calculated using the following loss function:

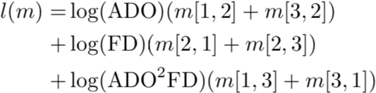

where *m* is a confusion matrix generated across all loci of a cell in which the first index represents the genotype that was measured for that particular cell (1 = homozygous reference, 2 = heterozygous, 3 = homozygous alternate), and the second index represents the genotype implied by the tree node. ADO = 0.01 and FD = 0.001 were used. Every cell was assigned to the node with the lowest loss *l*. For the trees presented in Extended Data Fig.2b-o and Extended Data Fig.9c-k only the numbers of cells with exact genotype matches were reported.

### Testing for evidence of homozygous genotypes

Due to the nature of our loci-specific mutation encoding (each gene is encoded as two loci), homozygous mutations are placed in the clonal hierarchy independently of their accuracy. Therefore, for every patient and at every locus with observed homozygous alternate genotype calls, the tested null hypothesis was that all homozygous alternate genotype calls are due to allelic dropout at a level not exceeding 0.05 using a one-tailed binomial test. The total number of draws for the test is the number of heterozygous and homozygous alternate genotype calls at the locus, the number of successful draws is the number of homozygous alternate calls, and the success rate is 0.05. Only homozygous alternate genotype calls below this 0.05 cut-off were reported in Extended Data Fig.2b-o and Extended Data Fig.9c-k; the results of the binomial test are reported for each patient and mutation in TableS8.

### Computational validation of *TP53* biallelic status from single-cell targeted genotyping datasets

To further validate the biallelic status of *TP53* mutations in our dataset, the patterns of allelic dropout in TARGET-seq single-cell genotyping data from patient carrying at least 2 different *TP53* mutations were investigated (n=6; Extended Data Fig.1j).

To test the hypothesis that the observed *TP53*-WT/*TP53*-homozygous (*TP53*-WT/HOM; or (0,2)) cells are the result of a chromosomal loss (and therefore, in different alleles), the following null hypothesis (H0) was formulated: observed *TP53*-WT/HOM cells are double allelic dropout events. Under H0, every *TP53*-WT/HOM cell (0,2), *TP53*-HOM/WT cell (2,0), *TP53*-HOM/HOM (2,2) as well as an unknown number of *TP53*-WT/WT (0,0) are the result of a *TP53*-HET/HET (1,1) cell undergoing allelic dropout (ADO) at both sites. The following assumptions were made: (a) ADO is unbiased towards HOM or WT and (b) ADO events at each *TP53* site are independent. The null hypothesis was then tested with a binomial test, where the number of (2,2) events should be half the sum of (0,2) + (2,0) events (Extended Data Fig.1j). (0,0) events were disregarded.

If *TP53* mutations are biallelic, the expected number of WT/HOM and HOM/WT would be higher than HOM/HOM cells taking into account TARGET-seq expected allelic dropout rates (1-5%).

### Single cell 3’-biased RNA-sequencing data pre-processing

FASTQ files for each single cell were generated using bcl2fastq (version 2.20) with default parameters and the following read configuration: Y8N*, I8, Y63N*. Read 1 corresponds to a cell-specific barcode, index read correspond to an i7 index sequence from each cDNA pool, and read 2 corresponds to the cDNA molecule. PolyA tails were trimmed from demultiplexed FASTQ files with TrimGalore (version 0.4.1). Reads were then aligned to the human genome (hg19) using STAR (version 2.4.2a) and counts for each gene were obtained with FeatureCounts (version 1.4.5-p1; options --primary). Counts were then normalized by dividing each gene count by the total library size of each cell and multiplying this value by the median library size of all cells processed, as implemented in the “*normalize_UMIs*” function from the SingCellaR package^67^ (https://github.com/supatt-lab/SingCellaR). A summary of the pre-processing pipeline can be found in https://github.com/albarmeira/TARGET-seq-WTA.

Quality control was performed using the following parameters: number of genes detected>500, percentage of ERCC derived reads<35%, percentage of mitochondrial reads<0.25%, percentage of unmapped reads<75%. Cells with less than 2000 reads in batch1, 5000 reads in batch2 and 10000 reads in batch3 were further excluded. This QC step was performed independently for each sequencing batch owing to differences in sequencing depth (mean library size: 42949 batch 1, 93580 batch 2 and 173145 batch3). After these QC steps, 7200 cells passed QC for batch1, 5838 for batch2 and 6490 for batch 3 (78.5%, 75.0% and 82.4% of cells processed, respectively). Then, 2733 cells from a previously published study^16^ corresponding to 8 myelofibrosis patients and 2 normal donor controls were further integrated, encompassing a final dataset of 22261 cells in total.

### Identification of highly variable genes

Highly variable genes above technical noise were identified by fitting a gamma generalized linear model (GLM) model of the log2(mean expression level) and coefficient of variation for each gene, using the “*get_variable_genes_by_fitting_GLM_model*” from SingCellaR package and the following options: *mean_expr_cutoff* = 1, *disp_zscore_cutoff* = 0.1, *quantile_genes_expr_for_fitting* = 0.6, *quantile_genes_cv2_for_fitting* = 0.2. Those genes with a coefficient of variation above the fitted model and expression cut-off were selected for further analysis, excluding those annotated as ribosomal or mitochondrial genes.

### CNA inference from single cell transcriptomes

InferCNV was used to identify CNAs in single-cell transcriptomes^68^ (https://github.com/broadinstitute/inferCNV/wiki). Briefly, inferCNV creates genomic bins from gene expression matrices and computes the average level of expression for each of these bins. The expression across each bin is then compared to a set of normal control cells, and CNAs are predicted using a hidden markov model. For each patient, inferCNV was performed with the following parameters: “*cutoff=0.1, denoise=T, HMM=T*”, compared to the same set of normal donor control cells (n=992). To identify CNA subclones, inferCNV in *analysis_mode=’subclusters’* was used. CNAs identified by inferCNV were manually curated by removing those with size<10kb, merging adjacent CNA calls with identical CNA status into larger CNA intervals and comparing them to SNP-Array bulk CNA calls. Finally, to generate combined TARGET-seq single-cell genotyping and CNA-based clonal hierarchies, the CNA status from each inferCNV cluster was assigned to its predominant genotype.

### Dimensionality reduction, data integration and clustering

PCA was performed using “*runPCA*” function from the *SingCellaR* R package, and Force-directed graph analysis was subsequently performed using the “*runFA2_ForceDirectedGraph*” with the top 30 PCA dimensions and the following options: *n.neighbors*=5, *fa2_n_iter*=1000 to generate the plots in Extended Data Fig.4a.

For the analysis of patient IF0131 presented in Extended Data Fig.3m, PCA was performed using “*runPCA*” function from the *SingCellaR* R package and then UMAP was performed using the “*runUMAP*” function with the top 10 PCA dimensions and the following options: *n.neighbors*=20, *uwot.metric* = “correlation”, *uwot.min.dist*=0.30, *n.seed* = 1.

Integration of TARGET-seq single-cell transcriptomes from 10538 cells corresponding to 14 *TP53*-sAML samples was performed using “*runHarmony*” function implemented in the SingCellaR package, using the patient ID as covariate and the following options: *n.dims.use*=20, *harmony.theta* = 1, *n.seed* = 1. Diffusion map analysis was performed using “*runDiffusionMap*” with the integrative Harmony embeddings and the following parameters: *n.dims.use*=20, *n.neighbors*=5, *distance*=”euclidean”. Signature scores were calculated using “*plot_diffusionmap_label_by_gene_set*” to generate the plots in Fig.2a and Fig.3a.

### Pseudotime trajectory analysis

Monocle3^69^ (https://cole-trapnell-lab.github.io/monocle3/) was used to infer differentiation trajectories from single cell transcriptomes. Raw UMI count matrix and clustering annotations were extracted from the SingCellaR object to build a Monocle3 ‘cds’ object. ‘*learn_graph’* function was then used calculate the trajectory, using *TP53*-WT preleukemic cell cluster as the root node. Pseudotime was calculated using ‘*order_cells’* function and overlayed on the diffusion map embeddings to generate the plot in Fig.2b.

### Differential expression analysis

Differentially expressed genes from TARGET-seq datasets were identified using a combination of non-parametric Wilcoxon test, to compare the expression values for each group, and Fisher’s exact test, to compare the frequency of expression for each group, as previously described^17^. Logged normalized counts were used as input for this comparison, including genes expressed in at least 2 cells. Combined p-values were calculated using Fisher’s method and adjusted p-values were derived using Benjamini & Hochberg procedure. Significance level was set at p-adjusted<0.05. For the analysis presented in Extended Data Fig.4b and TableS2, the top 100 differentially expressed genes with log2(fold-change)>0.3 and at least 20% expressing cells are shown. For the analysis presented in Fig.2k,l, only genes overexpressed in *TP53* multi-hit cells and log2(fold-change)>0.75 were included; for Fig.4c, only those with log2(fold-change)>1 were considered. Violin plots (Extended Data Fig.9l,m) from selected differentially expressed genes were generated using “ggplot2” package in R.

### Gene-Set Enrichment analysis

For analysis involving <500 cells per group (Fig.4c, TableS5) GSEA was performed using GSEA software (https:/www.gsea-msigdb.org/gsea/index.jsp) with default parameters and 1000 permutations on the phenotype, using log2(normalized counts).

For analysis involving >500 cells per group (Fig.3k and Extended Data Fig.4c), GSEA was performed with “*identifyGSEAPrerankedGene”* function from *SingCellaR* R package with default options. Briefly, differential expression analysis was performed between two cell populations using Wilcoxon rank sum test and the resulting p-values were adjusted for multiple testing using the Benjamini-Hochberg approach. Prior to the differential expression analysis, down-sampling was performed so that both cell populations had the same number of cells. Next, -log10(p-value) transformation was performed and the resulting p-values were multiplied by +1 or -1 if the corresponding log2FC was>0.1 or <- 0.1, respectively. The genelist was ranked using this statistic in ascending order and used as input for GSEA analysis using “*fgsea”* function from the *fgsea* R package with default options.

MSigDB HALLMARK v7.4 50-gene sets or previously published signatures (https://www.gsea-msigdb.org/gsea/msigdb/cards/GENTLES_LEUKEMIC_STEM_CELL_UP) were used for all analysis. Normalised enrichment scores (NES) were displayed in a heatmap using *pheatmap* R package. Gene sets with False Discovery Rate (FDR) q-value lower than 0.25 were considered significant.

### Projection of single cell transcriptomes

A previously published human haematopoietic atlas was downloaded from https://github.com/GreenleafLab/MPAL-Single-Cell-2019 and used as a normal haematopoietic reference to project *TP53*-sAML and *de novo* AML transcriptions using Latent Semantic Index Projection (LSI)^70^. Common genes to all datasets were selected and then, *TP53*-sAML or previously published *de novo* AML cells^25^ were projected using “*projectLSI’* function for the analysis presented in Fig.2c,d. A previously published human myelofibrosis atlas^71^ was used as a reference to project *TP53*-sAML multi-hit cells in the analysis presented in Extended Data Fig.5a,b, using previously defined force-directed graph embeddings.

### Velocyto analysis

Loom files were generated for each single cell using velocyto (v0.17.13) with options *-c* and *-U,* to indicate that each BAM represents an independent cell and reads are counted instead of molecules (UMIs), respectively^72^. The individual loom files were subsequently merged using the *combine* function from the *loompy* python module.

Healthy donors with at least 300 cells with RNA-sequencing data and patients with at least 300 cells consisting of >50 preleukemic (*TP53* wildtype) cells and > 50 *TP53* multi-hit cells were included for analysis. For each individual, Seurat object was created from the merged loom file and processed for downstream RNA-velocity analysis^73^. Specifically, for each patient, the spliced RNA counts were normalised using regularised negative binomial regression with the *SCTransform* function^74^. Next, linear dimension reduction was performed using *RunPCA* function and the first 30 principal components were further used to perform non-linear dimension reduction using the *RunUMAP* function. Ninety-six multiple rate kinetics (MURK) genes previously shown to possess coordinated step-change in transcription and hence violate the assumptions behind scVelo were removed^75^. The processed and MURK gene-filtered Seurat object was then saved as h5Seurat format using the *SaveH5Seurat* function and finally converted to h5ad format using the *Convert* function.

AnnData object was created from the h5ad file using the *scvelo* python module for RNA velocity analysis^76^. Highly variable genes were identified and the corresponding spliced and unspliced RNA counts were normalized and log2-transformed using the *scvelo.pp.filter_and_normalize* function. Next, the 1^st^ and 2^nd^ order moments were computed for velocity estimation using the *scvelo.pp.moments* function. The velocities (directionalities) were computed based on the stochastic model as defined in the *scvelo.t1.velocity* function, and the velocities was subsequently projected on the UMAP embeddings generated from Seurat above. Finally, the UMAP embeddings were annotated using the HSPC and erythroid lineage signature scores ^67^, and *TP53* mutation status. For each cell, the cell lineage signature score was computed using the average *SCTransform* expression values of the individual cell lineage genes.

### Analysis of bulk BeatAML and TCGA gene expression datasets

#### Data retrieval and pre-processing

Two publicly available AML cohorts with genetic mutation and RNA-sequencing data available were used to validate findings from our single-cell analysis, namely BeatAML^26^ and The Cancer Genome Atlas (TCGA)^27^. Gene expression values in FPKM (fragments per kilobase of transcript per million mapped reads) were retrieved from the National Cancer Institute (NIH) Genomic Data Commons (GDC)^77^. Gene expression values were then offset by 1 and log2-transformed. *TP53* point mutation status was retrieved from the cBio Cancer Genomics Portal (cBioPortal)^78^. Clinical data including survival data for BeatAML and TCGA was retrieved from the BeatAML data viewer (Vizome) and NIH GDC, respectively.

We selected samples from the BeatAML cohort with an AML diagnosis (540 *de novo* AML and 96 secondary AML) collected within 1 month of the patient’s enrolment in the study, and with both *TP53* mutation status and RNA-sequencing data available. For patients in which multiple samples were available, samples were collapsed to obtain patient-level data. Specifically, the mean gene expression value for each gene from multiple samples was used to represent patient-level gene expression value. Furthermore, patients with at least one sample with a *TP53* mutation were considered *TP53-*mutant. Analysis of *TP53* variant allele frequency and reported karyotypic abnormalities indicated that the vast majority of patients could be classified as “multi-hit”, and therefore patients were classified as *TP53-*mutant or WT without taking into account *TP53* allelic status. In total, 360 patients with *TP53* mutation status (329 *TP53*-WT and 31 *TP53*-mutant) and RNA-sequencing data available were included for analysis. Of these, 322 patients had concomitant survival data available (294 *TP53*-WT and 28 *TP53*-mutant).

The TCGA cohort consisted for 200 *de novo* AML patients represented by one sample each, out of which 151 patients had *TP53* mutation status (140 *TP53*-WT and 11 *TP53*-mutant) and RNA-sequencing data available, and were included for analysis. Of these, 132 patients had concomitant survival data available (124 *TP53*-WT and 8 *TP53*-mutant).

#### Cell lineage gene signature scores

For each sample, a given cell lineage gene signature score was computed as the mean expression values of the individual genes belonging to the cell lineage gene signature. Here, the gene signature scores for two cell lineages were computed, namely myeloid and erythroid populations. Two gene sets for each cell lineage were compiled. The first gene set was based on cell lineage markers previously reported in the literature whereas the second gene set was based on cell lineage markers derived from analysing a published single-cell dataset^70^. Genes from each score are described in TableS3.

For the former approach, six erythroid genes (*KLF1, GATA1, ZFPM1, GATA2, GYPA, TFRC;* Fig.2e, Extended Data Fig.5h) and seven myeloid genes (*FLI1, SFPI1, CEBPA, CEBPB, CD33, MPO, IRF8;* Fig.2f) were identified. For the latter approach, the expression values of erythroid and myeloid cell clusters were first compared separately against all other cell clusters using Wilcoxon ranked sum test. The erythroid cluster consisted of the early and late erythroid populations while the myeloid cluster consisted of granulocyte, monocyte, and dendritic cell populations. Erythroid and myeloid-specific gene signatures were defined as genes having FDR values < 0.05 and log2 fold change > 0.5 in >=20 and 17 comparisons, respectively. In total, 100 erythroid genes and 135 myeloid genes were identified from this single-cell dataset (TableS3), and were used to compute the scores presented in Extended Data Fig.5c-f.

### Prognostic signatures and Cox-regression survival models

#### Leukaemic stem cell (LSC) signature score

The 17-gene leukaemic stem cell (LSC17) gene set was retrieved from Ng *et al* ^31^. For each sample, the LSC17 score was defined as the linear combination of gene expression values weighted by their respective regression coefficients.

To identify *TP53*-sAML leukaemic stem cell signatures from our TARGET single-cell dataset, two different approaches were used. First, differentially expressed genes were identified as overexpressed in all Lin^-^CD34^+^ *TP53* multi-hit cells regardless of their transcriptional classification (“p53-all-cells”) versus myelofibrosis, healthy donor and *TP53*-WT preleukaemic cells; this gene-set consists of 30 genes (TableS4a). For the second approach, the same analysis was performed, but *TP53* multi-hit cells transcriptionally defined as leukaemic stem cells (falling in the leukaemic stem cell-like cluster, Fig.2a, middle) were specifically selected; this gene-set is comprised of 102 genes (“p53LSC”; TableS4a).

Next, lasso cox regression with 10-fold cross-validation implemented in the *glmnet* R package was used to identify p53-all-cells and p53-LSC genes that were associated with survival and to estimate their respective regression coefficients^79^. Specifically, Harrel’s concordance measure (C-index) was used to assess the performance of each fitted model during cross-validation. The best model was defined as the fitted model with the highest C-index. Subsequently, the coefficient for each gene estimated using the best model was used to compute the gene signature scores. Only genes with non-zero coefficient values were included in the final gene set. In total, 27 and 51 genes were retained from the p53-all-cells and p53-LSC gene sets, respectively. For each sample, the gene signature score for each gene set was defined as the linear combination of gene expression values weighted by their respective regression coefficient^31, 79^. The list of p53-LSC and p53-all-cells gene signatures is provided in TableS4b.

#### Survival analysis

For each gene expression signature, patients were first split using the median gene expression signature score. This resulted in two groups of patients, namely patients with high expression scores (greater than or equal to the median) and patients with low expression scores (lower than the median).

The Cox proportional hazards regression model implemented by the *survival* R package was fitted to estimate the hazard ratio associated with each feature. Log-rank test was used to test the differences between survival curves. The features analysed here were LSC17, p53-all-cells and p53-LSC signatures. Patients with low gene expression signature scores (below median) and patients with *TP53* wildtype status were specified as the reference groups in the model. Kaplan-Meier curves were plotted using the *survminer* R package to visualize the probability of survival and sample size at a respective time interval.

### In vitro assays

#### Short-term liquid culture experiments and interferon treatment

For short-term liquid culture differentiation experiments (Fig.3j, Extended Data Fig.7g,h), 1, 5 or 10 cells from different Lineage^-^CD34^+^ HSPC populations (HSC CD34^+^CD38^-^ CD45RA^-^CD90^+^, MPP CD34^+^CD38^-^CD45RA^-^CD90^-^, LMPP CD34^+^CD38^-^CD45RA^+^, more committed progenitors CD34^+^CD38^+^) were directly sorted into a 96-well tissue culture plate containing 100 μL of differentiation media: StemSpan (Catalog #09650, StemCell Technologies), 1% Penicillin+Streptomycin, 20 % BIT9500 (Cat# 9500, StemCell Technologies), 10 ng/mL SCF (Cat #300-07, Peprotech), 10 ng/mL FLT3L (Cat# 300-19, Peprotech), 10 ng/mL TPO (Cat# 300-18-10, Peprotech), 5 ng/mL IL3 (Cat # 200-03, Peprotech), 10 ng/mL G-CSF (Cat# 300-23, Peprotech), 10 ng/mL GM-CSF (Cat# 300- 03, Peprotech), 1 IU/mL EPO (Janssen, erythropoietin alpha, clinical grade) and 10 ng/mL IL6 (Cat# 200-06, Peprotech).

For differentiation experiments involving recombinant IFNγ (R&D Systems, 285-IF-100) and IFNα (rhIFN-alpha-2a, PBL Assay Science; 11100-1) treatment (Fig.4i), 100-500 Lin^-^ CD34^+^ cells were directly sorted into a 96-well tissue culture plate containing 50 μL of 2X differentiation media as described above, and incubated for 1 hour at 37°C 5% CO2. Then, an additional 50 μL of media containing 2X recombinant interferon was added to each well and mixed carefully, to generate a 1X IFNα dilution (final concentration 50 IU/μL) and 1X IFNγ dilution (final concentration 2 ng/μL).

For all liquid culture experiments, 50 μL of fresh 1X differentiation media was added at day 4. Readout was performed by flow cytometry after 12 days of culture using the antibodies detailed in TableS7.c (Panel D).

### Long-term culture initiating-cell (LTC-IC) assay

50 cells from each Lin^-^CD34^+^ population (HSC; MPP; LMPP; CD38+) and donor type (HD, MF, *TP53*-sAML) were sorted in triplicate. Cells were resuspended in 100 μL of myelocult (Stem Cell Technologies, #H5150) + Hydrocortisone (10^-6^M; Stem Cell Technologies, Cat#74142) and plated into an irradiated supportive stromal cell layer (5000 SI/SI cells and 5000 M2-10B4 cells per well) in a 96-well tissue-culture plate coated with Collagen type I (CORNING; Cat#354236).

Medium was changed weekly and after 6 weeks of culture, cells were washed in IMDM+20%FCS and plated into 1.4 mL of cytokine-rich methylcellulose (Methocult H4435, Stem Cell Technologies). Colonies were scored 14 days later under an inverted microscope, and each colony was classified according to its morphology as BFU-E (Burst-forming unit erythroid), CFU-G (granulocyte), CFU-GM (granulocyte-macrophage), CFU-M (macrophage) or CFU-GEMM (granulocyte, erythrocyte, macrophage, megakaryocyte). Selected colonies were used for cytospin and genotyping as outlined below.

### LTC-IC colony genotyping

LTC-IC colonies were picked from methylcellulose media, washed, resuspended in 10 μL of PBS and transferred to individual wells in a 96-well PCR plate. 15 μL of lysis buffer (Triton X-100 0.4%, Qiagen Protease 0.1 AU/mL) were added to each well and samples were incubated at 56 °C for 10 minutes and 72 °C for 20 minutes. A 3 μL aliquot from each lysate was used as input to generate a targeted and Illumina-compatible library for colony genotyping. The preparation of single cell genotyping libraries involves 3 PCR steps. In the first PCR step, target-specific primers spanning each mutation of interest are used for amplification (TableS6a); in the second PCR step, nested target-specific primers (TableS6b) attached to universal CS1 / CS2 adaptors (Forward adaptor, CS1: ACACTGACGACATGGTTCTACA; Reverse adaptor, CS2: TACGGTAGCAGAGACTTGGTCT) further enrich for target regions and in the third PCR step, Illumina-compatible adaptors containing sample-specific barcodes are used to generate sequencing libraries.

### Apoptosis experiments under IFNγ treatment

500 Lin^-^CD34^+^ cells were sorted into StemSpan (Catalog # 09650, StemCell Technologies) supplemented with 1% Penicillin+Streptomycin, 20 % BIT9500 (Cat# 9500, StemCell Technologies), 10 ng/mL SCF (Cat #300-07, Peprotech), 10 ng/mL FLT3L (Cat# 300-19, Peprotech), 10 ng/mL TPO (Cat# 300-18-10, Peprotech), 5 ng/mL IL3 (Cat # 200-03, Peprotech) and 2 ng/μL rhIFNγ (R&D Systems, 285-IF-100). Cell were incubated at 37 C 5% CO_2_ and 24 hours later, washed with AnnexinV Binding buffer 1X, stained with 1:100 AnnexinV-PE (Biolegend, Cat# 640907), DAPI and analysed immediately by flow cytometry.

### TP53 knockdown and differentiation of human CD34+ cells

shRNA sequence for p53 knockdown has been previously cloned into the lentiviral vector pRRLsin-PGK-eGFP-WPRE and validated^80^. Primary human CD34^+^ cells from patients with MPN (Table S1) were infected twice with scramble (shCTL) or shTP53 with a MOI (Multiplicity of Infection) of 15 and sorted 48h later on CD34 and GFP expression. Cells were cultured in serum-free medium with a cocktail of human recombinant cytokines containing EPO (1 U/mL, Amgen), FLT3-L (10 ng/mL, Celldex Therapeutics, Inc.), G-CSF (20 ng/mL, Pfizer), IL-6 (10 ng/mL, Miltenyi), GM-CSF (5 ng/mL, Peprotech), IL-3 (10 ng/mL, Miltenyi), TPO (10 ng/mL, Kirin Brewery) and SCF (25 ng/mL, Biovitrum AB).

At day 12 of culture, cells were stained with the antibodies detailed in TableS7.c, Panel C. DAPI was used for dead cell exclusion before acquisition on a FACSCanto II (BD Biosciences) instrument. Analysis of FACS data was performed using Kaluza (Beckman Coulter) software.

### Quantitative real time PCR in shRNA experiments

In p53 knockdown experiments, RNA from either CD34^+^ cells sorted after transduction or bulk cells at day 12 of culture was extracted using Direct-Zol RNA MicroPrep Kit (Zymo Research) and reverse transcription was performed with SuperScript Vilo cDNA Synthesis Kit (Invitrogen). Quantitative RT-PCR was performed on a 7500 Real-Time PCR Machine using SYBR-Green PCR Master Mix (Applied Biosystems). Expression levels were normalized to *PPIA* (housekeeping gene). Primers used are listed in TableS6c.

### Xenotransplantation

Purified CD34^+^ cells from AML patients were transplanted via retroorbital vein injection in sublethally irradiated (1.5Gy) NOD.CB17-*Prkdcscid IL2rgtm1*/Bcgen mice (B-NDG, Envigo). All experiments were approved by the French National Ethical Committee on Animal Care (n° 2020-007-23589). Blood cell counts were performed monthly by submandibular sampling of mice with blood chimerism assessed by flow cytometry using hCD34, hCD45 and mCD45 antibodies (TableS7.b). At sacrifice (27 weeks or 31 weeks post-transplant), human bone marrow HSPC fractions were sorted on an Influx Cell sorter (BD Biosciences) after staining with the antibodies detailed in TableS7.b.

### Evaluation of cell morphology

Cell morphology from PDX models (Extended Data Fig.3d) and *in vitro* LTC-IC cultures (Extended Data Fig.7e) was assessed after cytospin of 50-100,000 cells onto a glass slide (5 min at 500 rpm) and May-Grünwald Giemsa staining, according to standard protocols. Images were obtained using an AxioPhot microscope (Zeiss).

### Mouse Bone Marrow Chimaeras

*Trp53*^tm2Tyj^ *Commd10*^Tg(Vav1-icre)A2Kio^ (hereafter referred to as Trp53^R172H/+^) CD45.1 mice and CD45.2 wild-type mice used for BM chimera experiments and IFNγ ELISA assays were bred and maintained in accordance to UK Home Office regulations. All experiments carried out in the UK were performed under Project License P2FF90EE8 approved by the University of Oxford Animal Welfare and Ethical Review Body.*Trp53*^tm2Tyj 81^ and *Commd10*^Tg(Vav1-icre)A2Kio 82^ (Jackson laboratory stock number #008610) have been previously described.

1 million bone marrow (BM) cells from Trp53^R172H/+^ CD45.1 mice and 1 million BM CD45.2 wild-type competitor mice were transplanted intra-venously into lethally irradiated (10 Gy total body irradiation, split dose) congenic CD45.2 mice. In each cohort, a selection of mice were injected intra-peritoneally with 3 rounds of 6 injections each of 200μg poly(I:C) (GE Healthcare, #27-4732-01). Poly(I:C) was administered during weeks 6-7, 10-11, 14-15. Within each round, injections were spaced one or two days apart. Analysis of peripheral blood chimerism was performed every 4 weeks, while BM chimerism was analysed 20 weeks after transplantation. Chimerism was assessed by flow cytometry (using the antibodies detailed in TableS7.d. 7AAD (Sigma) was used for dead cell exclusion. FACS analyses were carried out on BD Fortessa or BD Fortessa X20 (BD Biosciences) and profiles were later analysed using FlowJo software (version 10.1, BD Biosciences).

### IFNγ ELISA assay

Wild-type mice were injected intra-peritoneally with a single dose of 200 μg poly(I:C) and spleens were collected from injected mice and non-treated controls 4 hours after injection. Spleens were processed into a single-cell suspension in 200 μl PBS, spun down at 500g for 5 minutes and supernatant was collected and used as spleen serum. IFNγ levels were assessed using mouse IFNγ Quantikine ELISA assay (R&D Systems, cat MIF00) following the manufacturer’s instructions. 450nm and 540nm optical densities were determined using Clariostar microplate reader (BMG Labtech).

### Statistical analysis

Statistical analyses are detailed in Figure Legends and performed using GraphPad Prism software (7 or later version) or R (version 3.6.1) software. Number of independent experiments, donors and replicates for each experiment are detailed in Figure Legends.

### Data and code availability

Scripts to reproduce all figures will be uploaded in GitHub (https://github.com/albarmeira/) upon publication. Raw sequencing data will be made available through GEO (GSEXXXXXX) and targeted single-cell genotyping data will be made publicly available through SRA (SRAXXXXXX).

## Acknowledgments

We are grateful to patients and donors; without their generosity, this study would not have been possible. We also thank Steve Knapper, clinical study teams and other investigators involved in supporting sample collection, and King’s Health Partners Biobank for providing access to samples. We thank William Vainchenker for his scientific input; Zemin Ren and Timothé Denaes for help with mouse experiments and Sally-Ann Clark for help with sorting. This work was funded by a Medical Research Council (MRC) Senior Clinical Fellowship (A.J.M.; MR/L006340/1), a CRUK Senior Cancer Research Fellowship (A.J.M.; C42639/A26988.), a Cancer Research UK (CRUK) DPhil Prize Studentship (C5255/A20936) to A.R-M, a British Spanish Society Scholarship to A.R-M., a MRC Confidence in Concept/MLSTF Grant to A.R-M. and A.J.M (MC_PC_19049), the MRC Molecular Haematology Unit core award (A.J.M and S.E.J. Eirik; MC_UU_12009/5), Emergence Cancéropôle Ile de France 2017 (I.A-D.), Association pour la Recherche contre le Cancer 2018 (I.A-D.), Siric-Socrate 2019 (I.A-D.), INCA-PLBIO 2020 (I.A-D.). A.L.C. was supported by Paris University (MENRT grant). The authors would like to acknowledge the flow cytometry facility at the MRC Weatherall Institute of Molecular Medicine (WIMM) which is supported by the MRC Human Immunology Unit; MRC Molecular Haematology Unit (MC_UU_12009); National Institute for Health Research (NIHR), Oxford Biomedical Research Centre (BRC); Kay Kendall Leukaemia Fund (KKL1057), John Fell Fund (131/030 and 101/517), the EPA fund (CF182 and CF170) and by the MRC WIMM Strategic Alliance awards G0902418 and MC_UU_12025. The authors acknowledge the contributions of Dr. Neil Ashley at the MRC Weatherall Institute of Molecular Medicine (WIMM) Single Cell Facility and MRC-funded Oxford Consortium for Single-Cell Biology (MR/M00919X/1). The authors would also like to acknowledge the contribution of the WIMM Sequencing Facility, supported by the MRC Human Immunology Unit and by the EPA fund (CF268), the Gustave Roussy flow cytometry platform and mouse facility. We also thank the Oxford Genomics Centre at the Wellcome Centre for Human Genetics (funded by Wellcome Trust grant reference 203141/Z/16/Z) for the generation and initial processing of the OmniExpress SNP array data. The results published here are in whole or part based upon data generated by the TCGA Research Network (https://www.cancer.gov/tcga) and the BeatAML team. The views expressed are those of the authors and not necessarily those of the National Health Service (NHS), the NIHR or the Department of Health.

## Author contributions

A.R.M. conceived the project, designed and performed experiments, performed computational analysis, analysed data and wrote the manuscript. R.N., A.L.C., H.R., J.O.S., E.L. and A.P. designed, performed experiments and analysed data, W.W.W., G.W. and W.W.K. performed computational analysis. J.E.M. collected primary samples and clinical and bibliographic data. C.D. provided clinical data. C.B. and M.B. analysed SNP-Array data. J.O.S., C.B., N.S., F.G., F.P., I.P., M.D., C.H. provided patients samples, clinical data and scientific input. C.M., H.G. analysed and provided patients and PDX biological data (NGS and SNP-array). A.H. performed and analysed patient’s targeted sequencing. S.E.J, B.P. and S.T provided scientific input and conceptualization. S.T. supervised computational analysis. I.A-D. conceived and supervised the project, analysed data and wrote the manuscript. A.J.M. conceived and supervised the project, provided clinical care and wrote the manuscript.

## Competing Interests statement

A patent relating to the TARGET-seq technique is licensed to Alethiomics Ltd, a spin out company form the University of Oxford with equity owned by B.P. and A.J.M. The other authors declare no competing interests.

## Materials & Correspondence

Requests for material(s) should be addressed and will be fulfilled by corresponding authors: Alba Rodriguez-Meira, Iléana Antony-Debré and Adam J. Mead.

## Supplementary Tables

**TableS1**. Clinical and genetic details from healthy donors and patients included in the study.

**TableS2.** Differentially expressed genes between *TP53* multi-hit HSPCs and *TP53*-WT cells.

**TableS3**. Genesets used to calculate gene expression signature scores in TARGET-seq and publicly available bulk-transcriptomic datasets.

**TableS4.** Differentially upregulated genes in *TP53* multi-hit cells (globally or LSCs) and genes selected by lasso regression to derive p53-all-cells and p53-LSC signatures.

**TableS5**. Gene signatures from *TP53* mutant heterozygous HSPCs from CP-*TP53*-MPN and pre-*TP53*-AML patients.

**TableS6.** Primers used throughout the experiments presented in the manuscript.

**TableS7**. Antibodies used for all experiments presented throughout the manuscript.

**TableS8.** Summary of mutation-specific homozygous status statistical testing for *TP53*-sAML and CP *TP53*-MPN patients. Related to Fig.1b-f; Fig.4b; Extended Data Fig.2b-o; Extended Data Fig.9c-k.

